# DI_/cle_, a Measure Consisting of Insulin Sensitivity, Secretion, and Clearance, Captures Diabetic States

**DOI:** 10.1101/2022.12.28.522156

**Authors:** Hikaru Sugimoto, Ken-ichi Hironaka, Tomoko Yamada, Kazuhiko Sakaguchi, Wataru Ogawa, Shinya Kuroda

## Abstract

**Context:** Insulin clearance is implicated in regulation of glucose homeostasis independently of insulin sensitivity and insulin secretion.

**Objective:** To understand the relation between blood glucose and insulin sensitivity, secretion, and clearance.

**Methods:** We performed a hyperglycemic clamp, a hyperinsulinemic-euglycemic clamp, and an oral glucose tolerance test (OGTT) in 47, 16, and 49 subjects with normal glucose tolerance (NGT), impaired glucose tolerance (IGT), and type 2 diabetes mellitus (T2DM), respectively. Mathematical analyses were retrospectively performed on this dataset.

**Results:** The disposition index (DI), defined as the product of insulin sensitivity and secretion, showed a weak correlation with blood glucose levels, especially in IGT (*r* = 0.04; 95% CI, −0.63– 0.44). However, an equation relating DI, insulin clearance, and blood glucose levels was well conserved regardless of the extent of glucose intolerance. As a measure of the effect of insulin, we developed an index, designated “DI/cle,” that is based on this equation and corresponds to DI divided by the square of insulin clearance. DI/cle was not impaired in IGT compared with NGT, possibly as a result of a decrease in insulin clearance in response to a reduction in DI, whereas it was impaired in T2DM relative to IGT. Moreover, DI/cle estimated from a hyperinsulinemic-euglycemic clamp, OGTT, or a fasting blood test were significantly correlated with that estimated from two clamp tests (*r* = 0.52; 95% CI, 0.37– 0.64, *r* = 0.43; 95% CI, 0.24– 0.58, and *r* = 0.54; 95% CI, 0.38– 0.68, respectively).

**Conclusions:** DI/cle can serve as a new indicator for the trajectory of changes in glucose tolerance.

## INTRODUCTION

The blood glucose concentration is maintained within a narrow range by feedback-compensatory responses in the body. Insulin sensitivity and insulin secretion are linked by such a relation, with insulin secretion increasing to maintain normal glucose homeostasis as insulin sensitivity declines (1). This compensatory response has been regarded as “moving up” the hyperbolic curve of the product of insulin sensitivity and insulin secretion, and the failure of this response, regarded as “falling off” the hyperbolic curve, is associated with the transition from normal glucose tolerance (NGT) to impaired glucose tolerance (IGT) and type 2 diabetes mellitus (T2DM) (1). The disposition index (DI), generally defined as the product of insulin sensitivity and insulin secretion, has therefore been adopted as a measure that correlates well with blood glucose levels (1,2).

Several models that explain this moving up–falling off phenomenon as an interplay among insulin sensitivity, insulin secretion, and pancreatic β-cell mass have been described (3–6). Such models account for the physiological features and pathological trajectory of glucose tolerance—including NGT, IGT, and T2DM—to a certain extent, but their conformity to clinical or experimental data is not high (6). Moreover, the moving up and falling off the curve of the product of insulin sensitivity and insulin secretion appear not to adequately represent the relation between blood glucose and insulin levels. For example, comparison of individuals with NGT at extremes of insulin sensitivity revealed that an approximately twofold increase in blood insulin levels was associated with an increase in insulin secretion of <50% (7,8), indicating that the increase in insulin secretion may not fully explain the increase in circulating insulin levels in individuals with insulin resistance (8).

Insulin clearance is implicated as a determinant of circulating insulin levels that is independent of insulin secretion (8–16). Insulin clearance likely contributes to the compensatory responses of the body, with a reduction in insulin clearance in the insulin-resistant state resulting in an increase in circulating insulin levels and a lessening of the demand for insulin secretion by β-cells (11). In some instances, changes in the circulating insulin levels do not reflect changes in insulin secretion, instead being primarily influenced by insulin clearance (17). Furthermore, changes in insulin clearance have been implicated in the pathogenesis of glucose intolerance (18–21). Whereas these various findings suggest the importance of insulin clearance in the physiology and pathology of glucose homeostasis, it is difficult to directly evaluate this parameter from clinical data (22).

Mathematical models provide insight into phenomena in which various factors mutually interact and for which effects of such factors cannot be measured directly (22–25). We previously developed a mathematical model based on temporal changes in circulating glucose and insulin levels during consecutive hyperglycemic and hyperinsulinemic-euglycemic clamps (23). Using this model, we estimated insulin clearance and characterized its role in the regulation of glucose homeostasis, and we identified a relation between insulin clearance and the product of insulin sensitivity and insulin secretion (23). However, this study did not examine the goodness-of-fit of the relation in individuals with each category of glucose tolerance.

The objectives of this study were (1) to test the hypothesis that the product of insulin sensitivity and insulin secretion cannot fully represent blood glucose levels and insulin clearance, (2) to derive a better equation relating these parameters from an empirical method that necessitates numerical calculations and simple mathematical models that can be solved analytically, and (3) to develop a simple index to assess the relation. We here propose a new index, “DI/cle,” that consists of DI corrected by insulin clearance. We have characterized DI/cle in individuals with various levels of glucose tolerance, and we provide evidence that this new index can serve as an indicator of the effect of insulin on glucose homeostasis and can capture the transition from IGT to T2DM.

## MATERIALS AND METHODS

### Subjects and Measurements

We analyzed a previously reported data set (23) in the present study. This study was approved by the ethics committee of Kobe University Graduate School of Medicine (UMIN000002359). Participants were recruited between October 2008 and December 2011. In brief, a hyperglycemic clamp was performed by intravenous infusion first of a bolus of glucose over 15 min and then of a variable amount of glucose to maintain the plasma glucose level at 200 mg/dL. Ten minutes after the end of the hyperglycemic clamp, a hyperinsulinemic-euglycemic clamp was performed by intravenous infusion of insulin with a target plasma glucose level of 90 mg/dL or of the fasting level in the case of NGT and IGT subjects whose fasting plasma glucose level was <90 mg/dL. Blood was obtained from arterialized veins that were confirmed by oxygen saturation. These clamp analyses and a 75-g oral glucose tolerance test (OGTT) were performed within a period of 10 days.

Fifty NGT, 18 IGT, and 53 T2DM subjects were initially used for parameter estimation. As we previously described (23), three NGT, two IGT, and three T2DM individuals were subsequently eliminated as outliers. In addition, one T2DM individual was eliminated because of missing values, with the remaining 47 NGT, 16 IGT, and 49 T2DM subjects being analyzed. Detailed characteristics of each subject are described in the previous study (23).

### Mathematical models

To investigate the relation among insulin sensitivity, insulin secretion, insulin clearance, and blood glucose levels, we examined a previously developed mathematical model (23), which is written as follows:

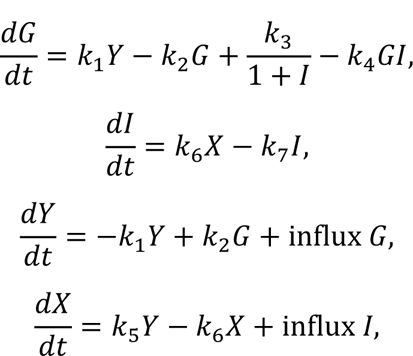

where *G* and *I* denote blood glucose and insulin concentrations, respectively. The variable *Y* corresponds to the effective glucose (the portion of glucose that affects insulin secretion) on induction of variable *X*, which can be regarded as insulin secreted from β-cells. The fluxes influx *G* and influx *I* denote glucose and insulin infusions, respectively. The indices *k*_4_, *k*_5_, and *k*_7_correspond to insulin sensitivity, insulin secretion, and insulin clearance, respectively. The parameters were estimated to minimize residual sum of the square (RSS) between the time course during clamp tests and the model trajectories, as previously described (23).

To validate the relation among the parameters, we also calculated insulin secretion, insulin sensitivity, and insulin clearance by using other methods. Insulin secretion was calculated from the deconvolution of the fasting serum C-peptide concentration (26). We investigated the first-phase insulin secretion, as follows:

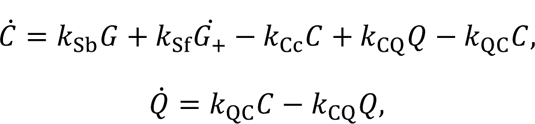

where *k*_Sb_, *k*_Sf_, and *k*_Cc_ correspond to basal insulin secretion, first-phase insulin secretion, and C-peptide clearance, respectively. The variable *C* and *Q* denote the concentrations of C-peptide in the accessible and non-accessible compartments, respectively (26). *k*_CQ_ and *k*_QC_ are rate parameters describing C-peptide exchange kinetics (26). By approximating the second-order derivative as zero and *k*_Sb_*G* = *k*_Cc_*C* at 0 min, we calculated *k*_Sf_ as follows:

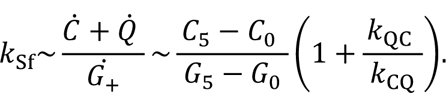

The subscripts of *C* and *G* denote the time (min) during a hyperglycemic clamp. Five individuals with *k*_Sf_ ≤ 0 were removed in this analysis.

As an index of insulin sensitivity, the insulin sensitivity index (ISI) was calculated, as described previously (23,27,28). As an index of insulin clearance, we calculated metabolic clearance rate of insulin (MCR) (23). To further examine the relation between blood insulin and glucose, we calculated basal insulin effect (29).

To further validate the relation among the parameters, and to analytically estimate the relation among the parameters, we also examined a simple and stable model, which can be written as follows (30):

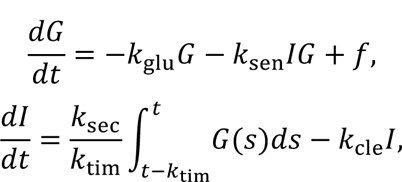

where *k*_glu_, *k*_sen_, *k*_sec_, *k*_tim_, and *k*_cle_ are glucose effectiveness, insulin sensitivity, insulin secretion, the duration of the preceding interval whose plasma glucose influence the current insulin secretion, and insulin clearance, respectively. *f* is the sum of the constant baseline hepatic glucose release and the amount of infused glucose.

To investigate how the relation among the parameters change with the progression of glucose intolerance, we also examined *β*GI model (5,19), which has been used to explore the characteristics of glucose homeostasis and is written as follows:

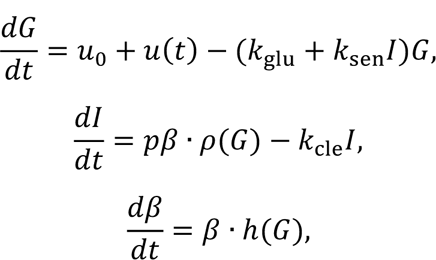

where *u*_0_, *u*(*t*), *p*, and *β* are endogenous glucose production and infused glucose, insulin secretion and beta-cell functional mass, respectively. *ρ*(*G*) is a monotonically increasing function of *G*. ℎ(*G*) is the beta-cell growth rate, and at the glucose set point, *G* = *G*_0_, ℎ(*G*_0_) = 0.

### Statistical Analysis

We first evaluated the equations relating insulin sensitivity, insulin secretion, insulin clearance, and blood glucose levels, which were given by the following equations:

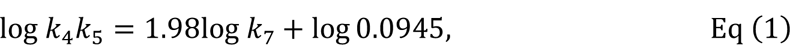

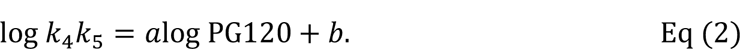

Eq (1) is a previously reported relation between insulin clearance and the product of insulin sensitivity and secretion (23). Eq (2) is a relation between the plasma glucose concentration at 120 min during the OGTT (PG120) and the product of insulin sensitivity and secretion. For estimating the parameters *a* and *b*, RSS between variables was minimized with the use of a nonlinear least squares technique, as described previously (23). After evaluating the goodness-of-fit of these equations, we examined the goodness-of-fit of various other equations to determine if a better equation existed. The goodness-of-fit was evaluated with Pearson’s correlation test (*r*). These values were reported with 95% confidence intervals (CIs) by using bootstrap confidence intervals. The number of resamples performed to form the bootstrap distribution was set at 10000. We also tested the relations among the variables by standardized major axis regression (31).

Comparisons among NGT, IGT, and T2DM individuals were performed by the Steel-Dwass test, a nonparametric test for multiple comparisons. A *P* value of <0.05 was considered statistically significant. The association between indices was evaluated with Spearman’s correlation test. The correlation coefficients were reported with 95% CIs by using bootstrap confidence intervals.

## RESULTS

### Derivation of a Conserved Equation for NGT, IGT, and T2DM from the Relations Among Insulin Sensitivity, Insulin Secretion, Insulin Clearance, and Blood Glucose Levels

We first investigated relations among insulin sensitivity, insulin secretion, insulin clearance, and blood glucose levels. We previously analyzed the relations using serum insulin and plasma glucose data obtained during consecutive hyperglycemic and hyperinsulinemic-euglycemic clamps in 47 NGT, 16 IGT, and 49 T2DM individuals (23), and we identified a relation regarding the three insulin-related parameters (23). This relation between insulin clearance and the product of insulin sensitivity and insulin secretion, was given by the following equation (23):

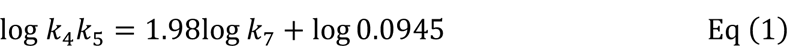

The indices *k*_4_, *k*_5_, and *k*_7_, which correspond to insulin sensitivity, insulin secretion, and insulin clearance, respectively, were derived from the previous study (23).

We reevaluated the goodness-of-fit of the relation in individuals with each category of glucose tolerance. The coefficient of determination (*R*^2^) of Eq (1) was estimated to be 0.26 for all subjects (Fig. S1A) and to be 0.28, 0.29, and –2.3 for NGT, IGT, and T2DM groups, respectively (Fig. S1B), indicating that the goodness-of-fit of Eq (1) was poor, especially for the T2DM group.

DI has been generally considered to reflect glucose disposal ability (2,32). The relation between DI and the blood glucose level can be represented as follows:

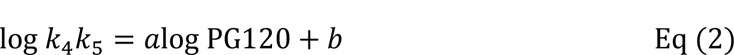

where the parameters *a* and *b* were estimated to minimize RSS between log *k*_4_*k*_5_ and *a*log PG120 + *b*, and PG120 indicates the plasma glucose concentration at 120 min during the OGTT, an important parameter for evaluation of glucose tolerance. Since the purpose of this study was to investigate the relation among insulin secretion, insulin sensitivity, insulin clearance, and blood glucose, DI was based on insulin secretion estimated by mathematical models rather than the insulin concentration, which is affected by both insulin secretion and clearance. Of note, the index defined in this way is also called adaptation index (33). The *r* value of Eq (2) estimated for all subjects was 0.81 (Fig. 1A). However, *r* of Eq (2) for the NGT, IGT, and T2DM groups was 0.41, 0.04, and 0.46, respectively (Fig. 1B), indicating that the goodness-of-fit was particularly poor for IGT. The parameters *a*_*n*_, *a*_*i*_, and *a*_*t*_ estimated from Eq (2) were –2.5, –0.24, and –1.8, respectively, and the parameters *b*_*n*_, *b*_*i*_, and *b*_*t*_ were 3.2, –1.7, and 1.2, respectively (Fig. 1B), suggesting that the relation between DI and blood glucose levels changes with the progression of glucose intolerance. Collectively, these results indicate that factors other than insulin secretion and insulin sensitivity may be involved in glucose dynamics.

**Figure 1.**
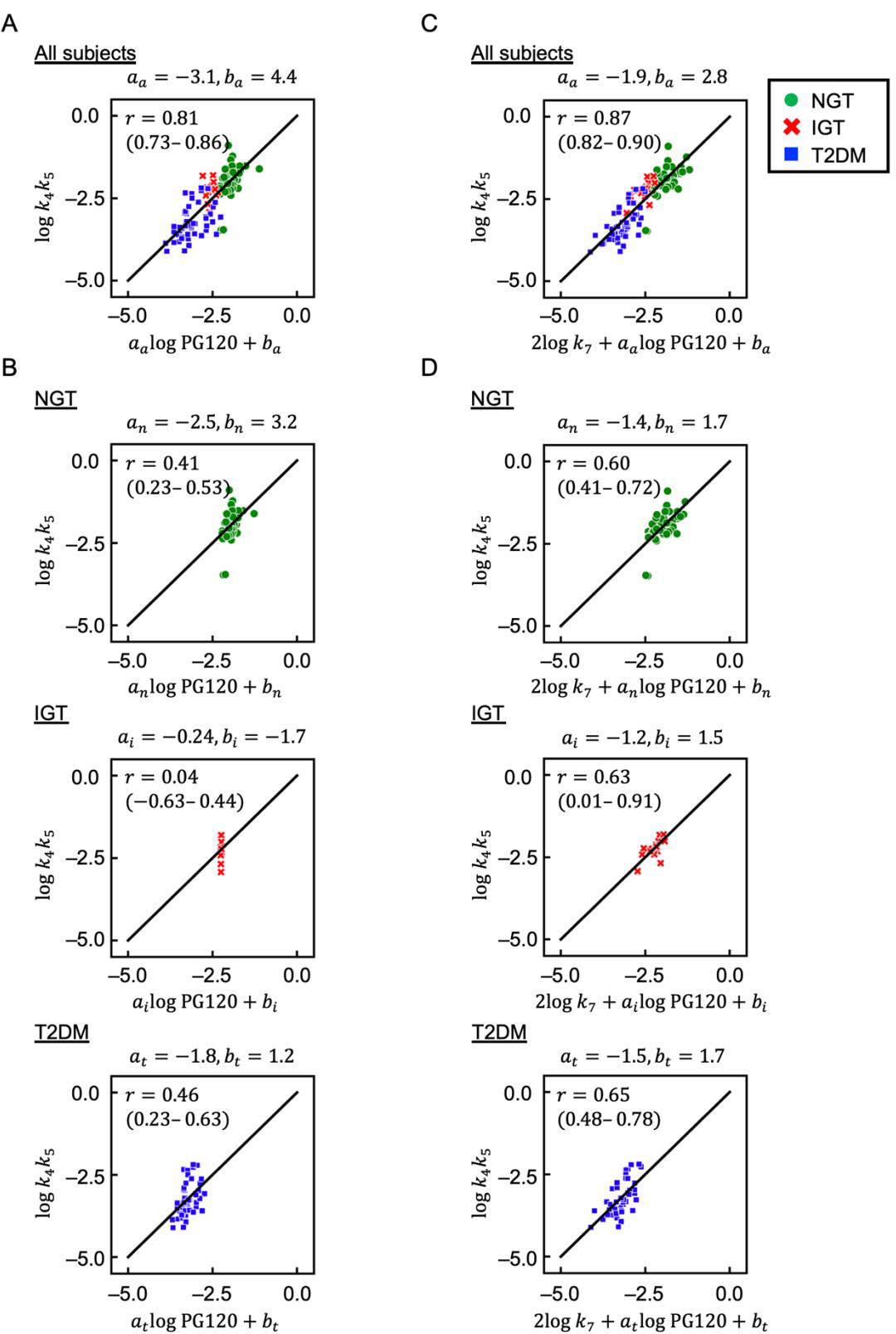
Relation between insulin-related parameters and blood glucose levels. (A) Scatter plot for *a*_*a*_log PG120 + *b*_*a*_ versus log *k*_4_*k*_5_ (Eq (2)). Each point corresponds to the values for a single subject. Green circles, red crosses, and blue squares indicate NGT, IGT, and T2DM subjects, respectively. *r* is Pearson’s correlation coefficient, and the value in the parenthesis is 95% confidence interval. The parameters *a*_*a*_ and *b*_*a*_ were estimated to minimize RSS between *a*_*a*_log PG120 + *b*_*a*_ and log *k*_4_*k*_5_ in all subjects. (B) Scatter plots for *a*log PG120 + *b* versus log *k*_4_*k*_5_ in NGT, IGT, and T2DM subjects (Eq (2)). The parameters *a*_*n*_ and *b*_*n*_, *a*_*i*_ and *b*_*i*_, and *a*_*t*_ and *b*_*t*_ were estimated to minimize RSS between *a*log PG120 + *b* and log *k*_4_*k*_5_ in NGT, IGT, and T2DM subjects, respectively. (C) Scatter plot for 2log *k*_7_ + *a*_*a*_log PG120 + *b*_*a*_ versus log *k*_4_*k*_5_ (Eq (3)). The parameters *a*_*a*_ and *b*_*a*_ were estimated to minimize RSS between 2log *k*_7_ + *a*_*a*_log PG120 + *b*_*a*_ and log *k*_4_*k*_5_ in all subjects. (D) Scatter plots for 2log *k*_7_ + *a*log PG120 + *b* versus log *k*_4_*k*_5_in NGT, IGT, and T2DM subjects (Eq (3)). The parameters *a*_*n*_ and *b*_*n*_, *a*_*i*_ and *b*_*i*_, and *a*_*t*_ and *b*_*t*_ were estimated to minimize RSS between 2log *k*_7_ + *a*log PG120 + *b* and log *k*_4_*k*_5_in NGT, IGT, and T2DM subjects, respectively. The indices *k*_4_, *k*_5_, and *k*_7_, which correspond to insulin sensitivity, insulin secretion, and insulin clearance, respectively, were calculated by a previously described method (23), and PG120 is the plasma glucose concentration at 120 min during an OGTT.

Given that the goodness-of-fit of Eq (1) and Eq (2) was poor, we attemted to show a better equation. Empirically, we hypothesized that the following equation might have a better fit:

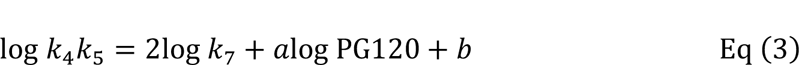

The parameters *a* and *b* were estimated to minimize RSS between log *k*_4_*k*_5_ and 2log *k*_7_ + *a*log PG120 + *b*. The *r* value of Eq (3) estimated for all subjects was 0.87 (Fig. 1C), which is better than that of Eq (2). The *r* value of Eq (3) for the NGT, IGT, and T2DM groups was 0.60, 0.63, and 0.65, respectively (Fig. 1D). The parameters *a*_*n*_, *a*_*i*_, and *a*_*t*_ estimated from Eq (3) were –1.4, –1.2, and –1.5, respectively, and the parameters *b*_*n*_, *b*_*i*_, and *b*_*t*_ were 1.7, 1.5, and 1.7, respectively (Fig. 1D). Differences in *r* and in the parameters of Eq (3) among NGT, IGT, and T2DM groups were smaller compared with those for Eq (2). These results indicated that Eq (3) in general provides a better representation of blood glucose levels than does Eq (2), especially for IGT. Of note, we here only indicated Eq (3) was one of the equations that have a better fit than Eqs (1) and (2), which have been considered to have good fit, and did not indicate that Eq (3) is the best. We further examined the characteristics of Eq (3) by different mathematical models and statistical methods, with the results confirming the validity of this equation (Appendix 1, Fig. S2, Fig. S3, and Fig. S4).

### “DI/cle”, a mearsure well correlated with blood glucose levels

Equation (3), which well describes the relation between the three insulin-related indices and blood glucose levels among NGT, IGT, and T2DM individuals, can be transformed as follows:

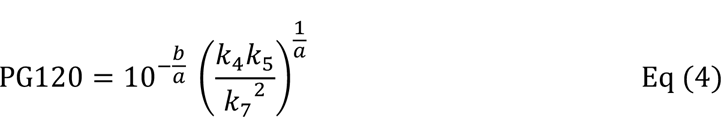

This transformation separates blood glucose concentration and the insulin-related indices onto the left and right sides. We designated this combination of the insulin-related indices on the right side (insulin sensitivity multiplied by insulin secretion and divided by the square of insulin clearance) as “DI/cle.” The product of *k*_4_and *k*_5_, which corresponds to DI, was significantly lower in individuals with IGT or T2DM than in those with NGT (23), whereas DI/cle was significantly lower in individuals with T2DM than in those with NGT or IGT but was similar in the latter two groups (Fig. 2A). These results were validated by the values of insulin sensitivity, insulin secretion, and insulin clearance estimated from the serum C-peptide concentration (Appendix 1 and Fig. S5). Moreover, DI/cle can be approximated as the extension of the basal insulin effect (29) to the insulin effect after glucose administration (Appendix 2). Of note, the dimension of DI/cle is dL/mg, which is the same as the dimension of the inverse of blood glucose levels. In addition, the sum of DI/cle and glucose effectiveness determines blood glucose levels after glucose administration (Appendix 3). Although the derivation from Eqs (1), (2), and (3) to Eq (4) is empirical and numerical, these analyses analytically yielded similar results regarding the relation between DI/cle and blood glucose levels, suggesting that DI/cle can literally be considered as the overall effect of insulin on blood glucose levels.

**Figure 2.**
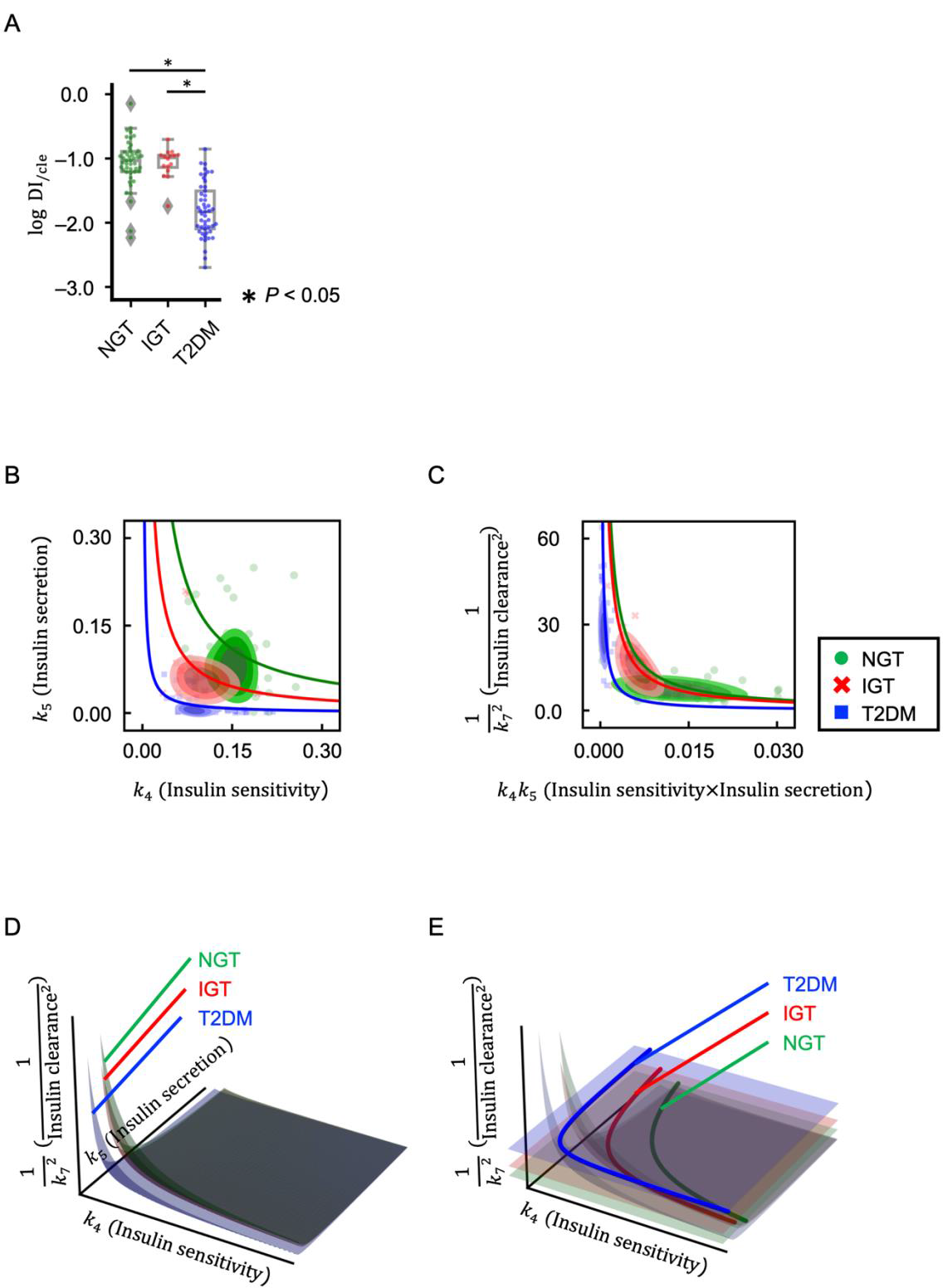
Relation between DI and DI/cle. (A) Box plots of log DI_/cle_, corresponding to the logarithm of *k*_4_*k*_5_/*k*_7_^2^, in NGT, IGT, and T2DM subjects. Each point corresponds to the value for a single subject. The boxes denote the median and upper and lower quartiles. *P* values were determined by the Steel-Dwass test. (B) Scatter plots for *k*_4_and *k*_5_fitted with hyperbolic curves (based on the function *xy* = constant), and kernel density plots. The hyperbolic curves correspond to the mean value of DI for NGT, IGT, or T2DM subjects. Each point corresponds to the values for a single subject, with green, red, and blue indicating NGT, IGT, and T2DM, respectively. The bandwidth of each kernel is based on the rule-of-thumb bandwidth. (C) Scatter plots for *k*_4_*k*_5_ and 1/*k*_7_^2^ fitted with hyperbolic curves (based on the function *xy* = constant), and kernel density plots. The hyperbolic curves correspond to the mean value of DI/cle in NGT, IGT, or T2DM subjects. Each point corresponds to the values for a single subject, with green, red, and blue indicating NGT, IGT, and T2DM, respectively. The bandwidth of each kernel is based on the rule-of-thumb bandwidth. (D) Hyperbolic surfaces (based on the function *xyz* = constant) corresponding to the mean value of DI/cle in NGT, IGT, or T2DM subjects. (E) Hyperbolic surfaces corresponding to the mean value of DI/cle as well as planes corresponding to the mean value of 1/insulin clearance^2^ in NGT, IGT, or T2DM subjects. The axes correspond to *k*_4_, *k*_5_, and 1⁄*k*_7_^2^ as in (D), but the label for *k*_5_ was omitted for clarity. The intersections (solid lines) of the surfaces representing DI/cle and the planes representing the mean value of 1/insulin clearance^2^ correspond to the hyperbolic curves in (B).

Given that DI is the product of insulin secretion and insulin sensitivity, it is generally represented as a hyperbolic curve with these two insulin-related parameters as the axes of coordinates. The hyperbolic curves corresponding to the mean values of DI in NGT, IGT, and T2DM subjects are shown in Figure 2B. On the other hand, DI/cle can be represented as a hyperbolic curve with DI and 1/insulin clearance^2^ as the axes of coordinates, given that DI/cle is the product of these two parameters. The hyperbolic curves corresponding to the mean values of DI/cle in NGT, IGT, and T2DM subjects are shown in Figure 2C. In general, as insulin sensitivity decreases, insulin secretion increases to maintain normal glucose homeostasis (the phenomenon of moving up the hyperbolic curve) (2). The relation between DI and 1/insulin clearance^2^ can also be interpreted as moving up the curve, with 1/insulin clearance^2^ increasing as DI decreases. Of note, we previously found that *k*_7_ (parameter for insulin clearance) was significantly smaller in individuals with IGT or T2DM than in those with NGT—that is, 1/insulin clearance^2^ increased with the progression of glucose intolerance (23).

We next investigated the relation between DI and DI/cle. We therefore represented DI/cle as a hyperbolic surface of insulin sensitivity, insulin secretion, and 1/insulin clearance^2^ (Fig. 2D and E). The mean values of DI/cle in NGT, IGT, and T2DM subjects that are shown as hyperbolic curves in Figure 2C are thus represented as hyperbolic surfaces in Figure 2D. Figure 2E shows the relation between Figure 2B and Figure 2D (that is, the relation between DI and DI/cle), with DI corresponding to the intersection of the surface representing DI/cle and the plane representing the mean value of 1/insulin clearance^2^. The DI for IGT subjects was significantly lower than that for NGT subjects (23), which can be interpreted as falling off the hyperbolic curve (Fig. 2B). DI/cle was similar for NGT and IGT (Fig. 2A), however, with the result that the corresponding hyperbolic curves (Fig. 2C) and hyperbolic surfaces (Fig. 2D) were almost identical. The difference in DI between NGT and IGT is reflected as a change in the intersection of the surface corresponding to DI/cle and the plane representing the mean value of 1/insulin clearance^2^ (Fig. 2E).

Of note, the phenomenon of moving up the curve of DI/cle for IGT represents almost the same phenomenon as predicted in a previous modeling study using the *β*GI model (19), indicating a pathway to failure of glucose homeostasis in which a change in insulin clearance may alter glucose dynamics even when the normal glucose set point is maintained (Appendix 4).

### Modified Mathematical Model of the Feedback Loop Linking Glucose and Insulin

We determined DI/cle by performing a hyperglycemic clamp and a hyperinsulinemic-euglycemic clamp to measure insulin secretion and insulin sensitivity together with insulin clearance (22,23,34), respectively. The performance of two clamp tests is laborious, however, and so we attempted to estimate DI/cle from a single clamp.

Given that we were not able to estimate DI/cle with a single clamp analysis using our previous model (23) (Appendix 5, Fig. S6, and Fig. S7A), we modified the model by changing the positions of influx *I* and influx *G* as shown in Figure 3A (Appendix 6). We simulated the time courses of blood glucose and insulin concentrations in the hyperinsulinemic-euglycemic clamp analysis, with one example for an NGT subject being shown in Figure 3B. To confirm that the modified model appropriately captured the characteristics of the time-series data, we evaluated the consistency between the ISI calculated from the actual measurements and that calculated from the simulation data. The coefficient of determination (*R*^2^) between measured ISI and simulated ISI was 0.96 (95% CI, 0.94 to 0.98) (Fig. 3C), indicating that the model was reasonably able to capture the essential characteristics of the time-series data.

**Figure 3.**
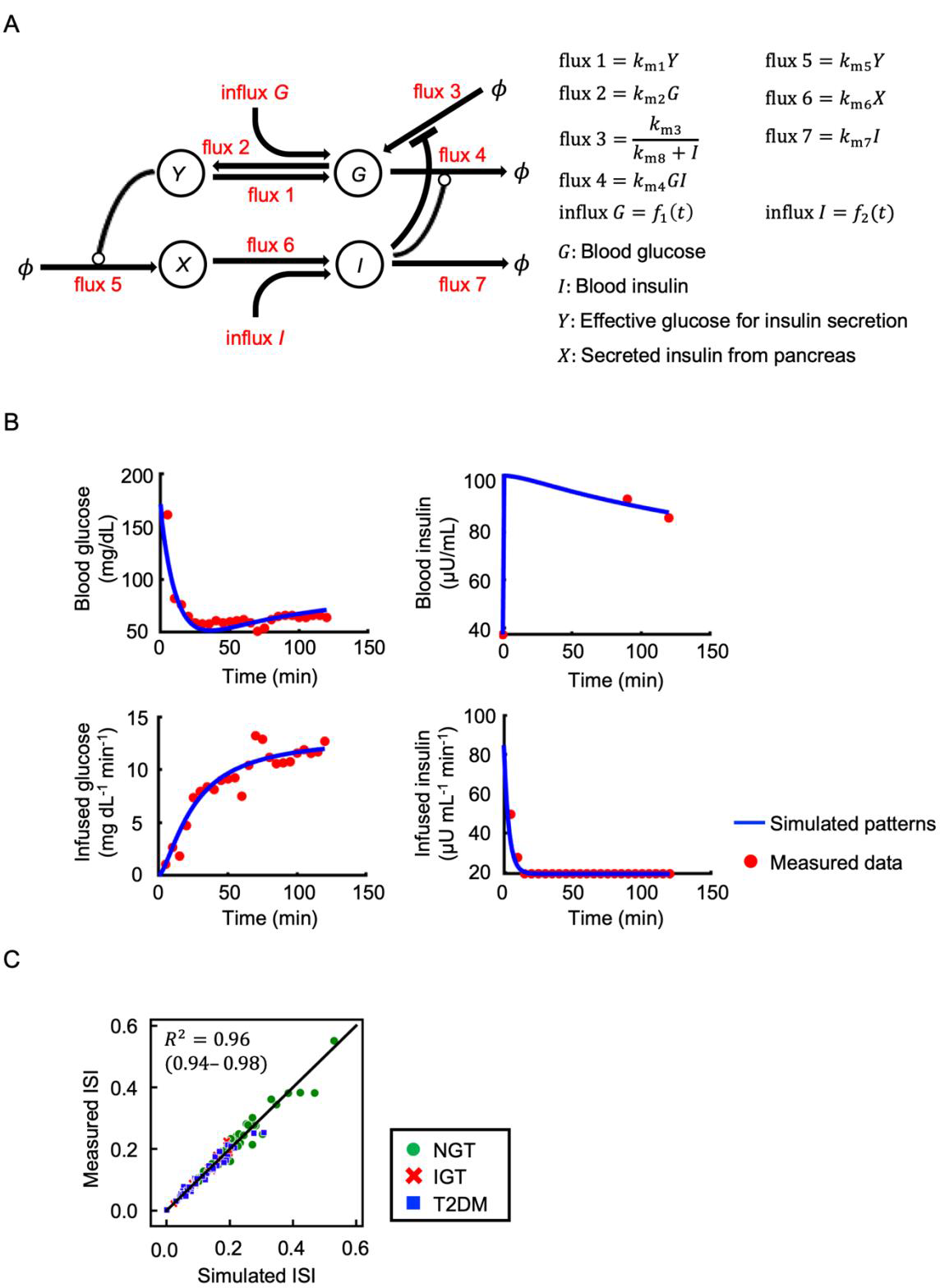
Modified mathematical model of the feedback loop that links glucose and insulin. (A) Model diagram. The letters within circles indicate the variables of the model, the arrows denote the fluxes, and the lines ending in circles or bars represent activation or inactivation, respectively. *ϕ* indicates a fixed value. The variables *G* and *I* represent blood glucose and insulin concentrations, respectively. The variable *Y* corresponds to the effective glucose concentration for induction of variable *X*, which represents insulin secreted from pancreatic β-cells. (B) Time courses of blood glucose, blood insulin, infused glucose, and infused insulin during hyperinsulinemic-euglycemic clamp analysis for an NGT subject. The blue lines indicate the simulated patterns, and the red circles indicate the measured time course data. The simulated values of each variable were rescaled to absolute concentration and plotted. (C) Scatter plot for simulated ISI versus measured ISI. Each point corresponds to the values for a single subject, with green circles, red crosses, and blue squares indicating NGT, IGT, and T2DM subjects, respectively. The measured and simulated ISI values were calculated from the normalized time course and plotted. *R*^2^ is the coefficient of determination, and the value in the parenthesis is 95% confidence interval.

### Prediction of DI/cle from only a hyperinsulinemic-euglycemic clamp

Some of the estimated parameters—including *k*_m7__ins, corresponding to insulin clearance estimated from the modified model for the hyperinsulinemic-euglycemic clamp alone— showed a tendency to diverge (Fig. 4A). However, *k*_m5_ /*k*_m7_^2^_ins was significantly correlated with *k*_5_/*k*_7_^2^_full (*r* = 0.36; 95% CI, 0.16 to 0.51), and *k*_m4_ *k*_m5_/*k*_m7_^2^_ins was significantly correlated with *k*_4_*k*_5_/*k*_7_^2^_full (*r* = 0.52; 95% CI, 0.37 to 0.64) (Fig. 4B and Fig. S7B). Of note, insulin sensitivity, insulin secretion, and insulin clearance estimated from the modified model for both the hyperglycemic clamp and the hyperinsulinemic-euglycemic clamp (*k*_m4_ _full, *k*_m5__full, and *k*_m7__full, respectively) were significantly correlated with those estimated from the original model (*k*_4__full, *k*_5__full, and *k*_7__full, respectively) (Fig. S8), indicating that both models can be used to estimate the insulin-related parameters when both the hyperglycemic and hyperinsulinemic-euglycemic clamps are performed. Thus, even with this modified model, Eq (3) showed a better fit than Eq (2): the *r* value of Eq (2) for the NGT, IGT, and T2DM groups was 0.03 (95% CI, –0.26 to 0.47), 0.25 (95% CI, –0.38 to 0.61), and 0.49 (95% CI, 0.24 to 0.66), respectively, and the *r* value of Eq (3) for the NGT, IGT, and T2DM groups was 0.97 (95% CI, 0.83 to 0.99), 0.81 (95% CI, 0.34 to 0.94), and 0.66 (95% CI, 0.46 to 0.79), respectively.

**Figure 4.**
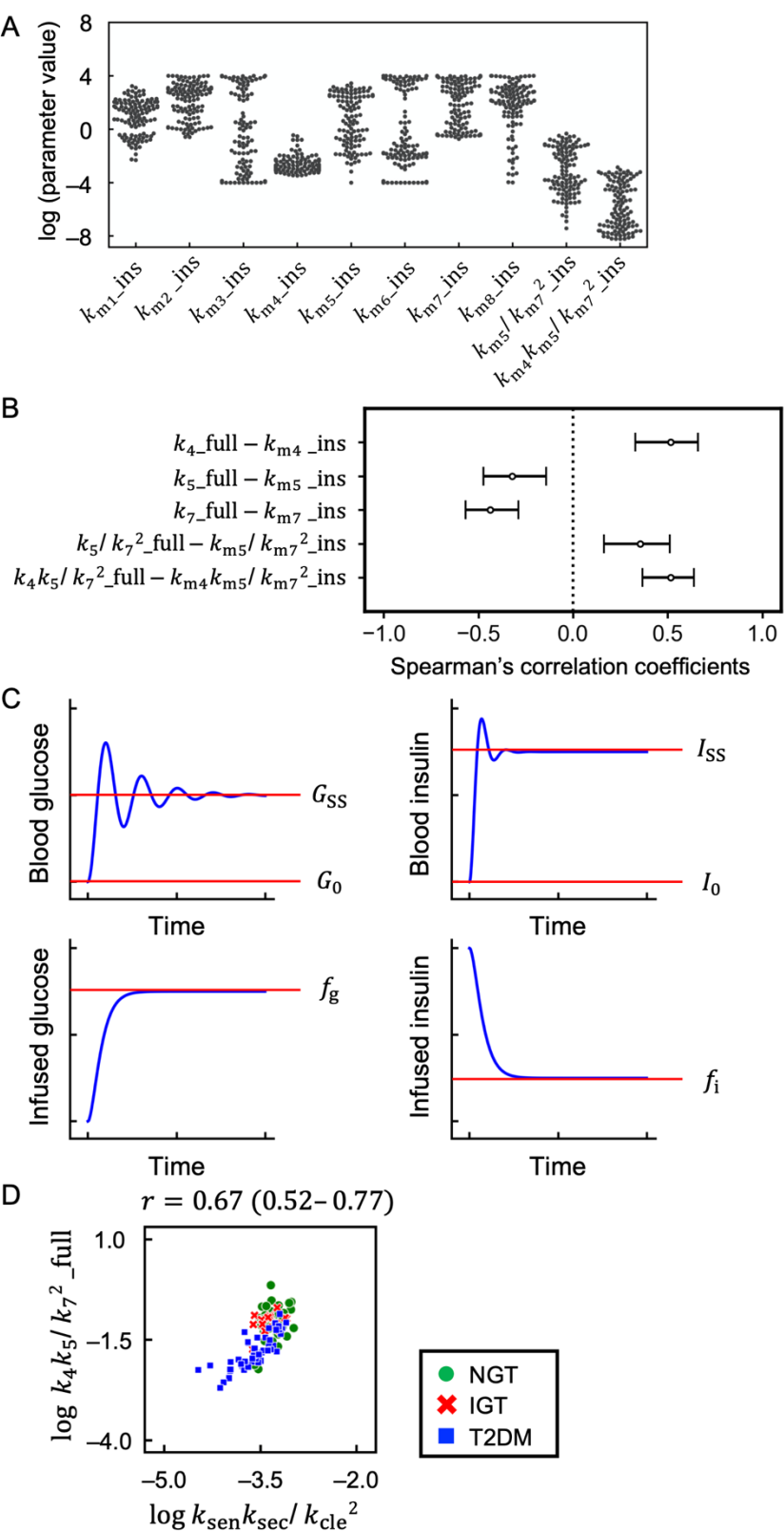
Relation between parameters estimated from a hyperinsulinemic-euglycemic clamp alone and those estimated from both hyperglycemic and hyperinsulinemic-euglycemic clamps. (A) Estimated parameters of the modified model based on a hyperinsulinemic-euglycemic clamp alone. (B) Spearman’s correlation coefficients. Bars indicate 95% confidence intervals. The parameters *k*_4__full, *k*_5__full, *k*_7__full, *k*_5_/*k*_7_^2^_full, and *k*_4_*k*_5_/*k*_7_^2^_full correspond to insulin sensitivity, insulin secretion, insulin clearance, the ratio of insulin secretion to the square of insulin clearance, and DI/cle estimated from both hyperglycemic and hyperinsulinemic-euglycemic clamps by the previously described mathematical model (23) (Fig. S6A), respectively. The parameters *k*_m4_ _ins, *k*_m5__ins, *k*_m7__ins, *k*_m5_ /*k*_m7_^2^_ins, and *k*_m4_ *k*_m5_/*k*_m7_^2^_ins correspond to insulin sensitivity, insulin secretion, insulin clearance, the ratio of insulin secretion to the square of insulin clearance, and DI/cle estimated from the hyperinsulinemic-euglycemic clamp alone by the modified mathematical model (Fig. 3A), respectively. (C) Simulated time courses of blood glucose, blood insulin, infused glucose, and infused insulin during a hyperinsulinemic-euglycemic clamp. *G*_SS_, blood glucose level at the steady state during insulin administration; *G*_0_, blood glucose level in the fasting state; *I*_SS_, blood insulin level at the steady state during insulin administration; *I*_0_, blood insulin level in the fasting state; *f*_g_, glucose infusion rate at the steady state; and *f*_i_, insulin infusion rate at the steady state. (D) Scatter plot for log *k*_sen_*k*_sec_/*k*_cle_^2^ versus log *k*_4_*k*_5_/*k*_7_^2^_full. The parameters *k*_sen_*k*_sec_/*k*_cle_^2^ and *k*_4_*k*_5_/*k*_7_^2^_full correspond to DI/cle analytically estimated from the hyperinsulinemic-euglycemic clamp alone (Appendix 7) and that estimated from the hyperglycemic and hyperinsulinemic-euglycemic clamps by the previously described mathematical model (23) (Fig. S6A), respectively. Each point corresponds to the values for a single subject, with green circles, red crosses, and blue squares indicating NGT, IGT, and T2DM subjects, respectively. *r* is Spearman’s correlation coefficient, and the value in the parenthesis is 95% confidence interval.

We also attempted to estimate DI/cle using a simple analytically computable method. We hypothesized that DI/cle can be analytically estimated as follows (Appendix 7):

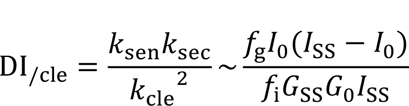

where *k*_sen_, *k*_sec_, and *k*_cle_ represent insulin sensitivity, insulin secretion, and insulin clearance, respectively, and *G*_SS_, *G*_0_, *I*_SS_, *I*_0_, *f*_g_, and *f*_i_ represent blood glucose level at the steady state during insulin administration, blood glucose level in the fasting state, blood insulin level at the steady state during insulin administration, blood insulin level in the fasting state, glucose infusion rate at the steady state, and insulin infusion rate at the steady state, respectively (Fig. 4C). This analytically calculated DI/cle (*k*_sen_*k*_sec_/*k*_cle_^2^) was significantly correlated with *k*_4_*k*_5_/*k*_7_^2^_full (*r* = 0.67; 95% CI, 0.52 to 0.77) (Fig. 4D). Of note, *k*_4_*k*_5_/*k*_7_^2^_full was also significantly correlated with DI/cle analytically estimated from a hyperinsulinemic-euglycemic clamp alone by using the serum C-peptide concentration (*r* = 0.71; 95% CI, 0.58 to 0.80), OGTT (*r* = 0.43; 95% CI, 0.24 to 0.58), or a fasting blood test (*r* = 0.54; 95% CI, 0.38 to 0.68) (Appendix 1, Appendix 8, Fig. S9 and Fig. S10).

## DISCUSSION

We have here derived an equation relating insulin sensitivity, insulin secretion, insulin clearance, and blood glucose levels [Eq (3)] that was well conserved among individuals with various levels of glucose tolerance, and we showed that this equation provides a better fit, particularly for individuals with IGT, than does an equation relating only insulin sensitivity, insulin secretion, and blood glucose levels [Eq (2)]. Insulin clearance is implicated in the regulation of blood glucose concentration (8), and our present data also suggest that, together with insulin sensitivity and insulin secretion, insulin clearance is important for such regulation, especially in individuals with IGT. Our present results also suggest that DI/cle, an index that corrects DI for insulin clearance, reflects glucose disposal ability, possibly by representing the effect of insulin on blood glucose concentration.

DI/cle is represented as a hyperbolic curve of DI and 1/insulin clearance^2^ or as a hyperbolic surface of insulin sensitivity, insulin secretion, and 1/insulin clearance^2^. The deterioration of glucose tolerance is sometimes explained by the failure of insulin secretion to increase sufficiently in compensation for a decline in insulin sensitivity, with this failure being referred to as falling off the hyperbolic curve of DI (2,32). Our data now suggest that, even when DI decreases, DI/cle is preserved if insulin clearance is also reduced. In general, moving up the hyperbolic curve of DI eventually positions individuals at unstable extreme points of the curve where the risk of disease onset increases (32), a phenomenon that has been referred to as canalization and decanalization (2,32) or as glucose allostasis (35). Given that individuals with IGT are at high risk of developing T2DM, the progression from NGT to IGT and T2DM can be referred to as canalization and decanalization of the hyperbolic surface of DI/cle.

Our data also indicate that IGT encompasses the condition of decreased insulin clearance with a relatively well preserved DI/cle and glucose set point (Appendix 4), and that T2DM encompasses that of decreased DI/cle with a disrupted glucose set point. A modeling study with simulated data indicated that changes in insulin clearance may affect glucose dynamics even when the normal glucose set point is maintained, and can constitute a pathway for progression of glucose intolerance (19). Our study based on mathematical models and observational data now indicates that the phenomenon predicted by this previous model can occur in IGT. Our findings are also consistent with those of a previous study indicating that an increase in insulin secretion is followed by a decrease in first-pass hepatic insulin clearance when insulin resistance is induced (15).

Despite the apparent analogy of the moving up–falling off phenomenon, whether the body possesses the ability to reduce insulin clearance in response to a decline in DI and the potential physiological mechanism underlying such a response remain unknown. However, evidence suggests that changes in insulin clearance can be a determinant of glucose intolerance (21,36). Whereas the direction of causation of decreased insulin clearance and glucose intolerance remains unclear, a classification based on DI/cle may provide new insight into the trajectories of glucose intolerance and T2DM.

Although these results were obtained from a limited number of individuals, the results were validated by several different mathematical models. Moreover, we attempted to estimate DI/cle from a hyperinsulinemic-euglycemic clamp alone, OGTT, or a fasting blood test so that the characteristics of DI/cle can be validated more easily in the future. Whereas we showed that DI/cle estimated from a hyperinsulinemic-euglycemic clamp test alone was significantly correlated with that estimated from the two clamps, performance of even a single clamp is labor-intensive. Moreover, the correlation between DI/cle estimated from the two clamps and that estimated from the single clamp was only 0.52, 0.67 or 0.71. It will therefore be important to develop more accurate and convenient methods for estimation of DI/cle in order to validate its physiological relevance and clinical usefulness.

The current study has several limitations. For simplicity, we considered only insulin sensitivity, insulin secretion, and insulin clearance. However, insulin secretion has basal, first-phase, and second-phase components (37), and both insulin sensitivity and insulin clearance have hepatic and peripheral components (3,22). Saturation of or time-dependent changes in insulin clearance (6,22) were not considered. Although the models used in this study possesses sufficient expressivity to fit the data from clamp tests (23) and IVGTT (26,30), and to conclude the significance of incorporating insulin clearance in addition to insulin secretion and insulin sensitivity, as well as a novel perspective on the progression of glucose intolerance with the *β*GI model, it is important to devise a more comprehensive measure that encompasses these more intricate effects in the future. Moreover, several better approaches to evaluation of the hyperbolic relation have also been presented, and a power function describing the relation between insulin sensitivity and insulin secretion gives a better representation (38–40). Given that insulin sensitivity and insulin clearance are correlated (41), the plane representing 1/insulin clearance^2^ is not necessarily horizontal (Fig. 2E), and the curve corresponding to DI also departs from a hyperbola when the plane departs from the horizontal. Therefore, this study indicates that insulin clearance should be considered when assessing the coefficient of disposition index, and future research on the coefficients of insulin sensitivity, insulin secretion, and insulin clearance in the formula defining the effect of insulin on blood glucose levels is warranted. In addition, the capacity for glucose disposal is affected not only by insulin, but also by other hormones including glucagon as well as by glucose (42,43). It is possible that differences in such factors may explain the glucose dynamics of glucose tolerance. Further studies are also warranted to develop extended equations that include glucose and glucagon to represent mechanisms of blood glucose regulation.

In conclusion, we have here shown that DI/cle, a new index that corrects DI by insulin clearance, better reflects changes in glycemia compared with DI. By analogy with falling off the hyperbolic curve of DI, the progression of glucose intolerance may be considered as falling off the hyperbolic surface of DI/cle. In theory, DI/cle represents the effect of insulin on blood glucose levels, and it may serve as an indicator for the trajectory of changes in glucose tolerance. Further study is warranted to understand the physiological and pathological relevance of this new index.

## Acknowledgments

We thank our laboratory members for critically reading this manuscript

## Contribution statement

H.S. and K.H. analyzed the data. H.S., K.H., T.Y., K.S., W.O., and S.K. wrote the manuscript. W.O. and S.K. supervised the study. S.K. is the guarantor of this work and, as such, had full access to all the data in the study and takes responsibility for the integrity of the data and the accuracy of the data analysis.

## Funding and Assistance

This study was supported by the Japan Society for the Promotion of Science (JSPS) KAKENHI (JP21H04759), CREST, the Japan Science and Technology Agency (JST) (JPMJCR2123), and The Uehara Memorial Foundation.

## Data availability

The data set analyzed in the present study was described in our previous study (23).

## Prior Presentation

A non–peer-reviewed version of this article was submitted to the bioRxiv preprint server (https://doi.org/10.1101/2022.12.28.522156) on 30 December 2022.

