## Supplementary file for "DI_/cle_, a Measure Consisting of Insulin Sensitivity, Secretion, and Clearance, Captures Diabetic States"

### Appendix

#### 1. Validity of Eq (3), the equation relating insulin sensitivity, insulin secretion, insulin clearance, and blood glucose levels

To further validate Eq (3), we investigated the equations in which the coefficient of $k_7$ is 1 or 3 (Fig. S2). The $r$ value of Eq (3) for the NGT, IGT, and T2DM groups was better than that of the equations in which the coefficient of $k_7$ is 1 or 3. Differences in the parameters of Eq (3) among NGT, IGT, and T2DM groups were smaller compared with those for the equations in which the coefficient of $k_7$ is 1 or 3 (Fig. S2), suggesting that Eq (3), whose coefficient of $k_7$ is 2, provides the best representation of blood glucose levels. To further investigate the goodness-of-fit of the equations in individuals with each category of glucose tolerance, we also conducted standardized major axis tests (Fig. S3), which have been used to investigate relationships among variables (1,2). The estimated coefficients of Eq (2) were statistically significantly different among NGT, IGT, and T2DM groups (Fig. S3A), but the differences in the estimated coefficients of Eq (3) were relatively small (Fig, S3B), indicating that Eq (3) provides a better representation of blood glucose levels than does Eq (2) for NGT, IGT, and T2DM groups.

We also estimated insulin secretion from the serum C-peptide concentrations, as previously described (3). As an index of insulin secretion ($k_{\mathrm{sec}}$), we calculated the insulin secretion rate at the fasting state (3) divided by the fasting glucose level. As an index of insulin sensitivity ($k_{\mathrm{sen}}$), we calculated insulin sensitivity index (ISI) (4). As an index of insulin clearance ($k_{\mathrm{cle}}$), we calculated metabolic clearance rate (MCR) (4,5). Of note, these parameters were estimated from a hyperinsulinemic-euglycemic clamp alone. Even when insulin sensitivity, insulin secretion, and insulin clearance were estimated in this way, Eq (3) provided a better representation of blood glucose levels than does Eq (2) (Fig. S4).

Instead of $k_{\mathrm{sec}}$, we also investigated the first-phase insulin secretion ($k_{\mathrm{Sf}}$). Even with this first-phase insulin secretion, Eq (3) tended to show a better fit than Eq (2): the $r$ value of

Eq (2) for the NGT, IGT, and T2DM groups was 0.06 (95% CI, –0.17 to 0.30), 0.08 (95% CI, –0.49 to 0.47), and 0.42 (95% CI, 0.02 to 0.61), respectively, and the $r$ value of Eq (3) for the NGT, IGT, and T2DM groups was 0.53 (95% CI, 0.28 to 0.71), 0.38 (95% CI, –0.08 to 0.71), and 0.35 (95% CI, 0.09 to 0.58), respectively. However, when insulin secretion was calculated in this way, the equation in which the coefficient of insulin clearance is 1 also showed a good fit: the $r$ value of the equation in which the coefficient of insulin clearance is 1 for the NGT, IGT, and T2DM groups was 0.53 (95% CI, 0.28 to 0.71), 0.37 (95% CI, –0.11 to 0.73), and 0.41 (95% CI, 0.11 to 0.61), respectively. This result may be because the dimension of insulin secretion differs depending on the calculation methods, as we discussed in Appendix 2. Either way, these results indicate the need to consider not only insulin secretion and insulin sensitivity but also insulin clearance to represent blood glucose levels, especially in NGT and IGT groups.

**2. Relation between "$DI_{/cle}$" and "basal insulin effect"**

This appendix details how "$DI_{/cle}$" can be approximated as the extension of basal insulin effect (BIE) (6) to the insulin effect during glucose administration. BIE is the effect of basal insulin on glucose uptake, and defined as follows (6):

$$\mathrm{BIE} = I_{\mathrm{basal}} k_{\mathrm{sen}}, \qquad \text{Eq (5)}$$

where $I_{\mathrm{basal}}$ and $k_{\mathrm{sen}}$ are the basal insulin concentration and insulin sensitivity, respectively. We extended the concept of BIE to a steady state insulin effect (SIE) during glucose administration as follows:

$$\mathrm{SIE} = I_{\mathrm{SS}} k_{\mathrm{sen}}, \qquad \text{Eq (6)}$$

where $I_{\mathrm{SS}}$ is the insulin concentration at the steady state. The minimal model, a mostly used mathematical model of glucose, cannot admit an equilibrium and, in some conditions, the concentration of insulin increases without bounds (7). Using a modified simple, stable, and unified model (7), $I_{\mathrm{SS}}$ can be transformed as follows:

$$I_{SS} = G_{SS}\frac{k_{sec}}{k_{cle}}, \quad \text{Eq (7)}$$

where $G_{SS}$, $k_{sec}$, and $k_{cle}$ are the glucose concentration at the steady state, insulin secretion, and insulin clearance, respectively.

Blood glucose levels after glucose administration ($G$) are experimentally approximated only by insulin clearance, as follows (4):

$$G \propto \frac{1}{k_{cle}}. \quad \text{Eq (8)}$$

Eq (6), (7), and (8) can be transformed and approximated as follows:

$$\text{SIE} \propto \frac{k_{sen}k_{sec}}{{k_{cle}}^2}. \quad \text{Eq (9)}$$

The right side of Eq (9) is the same as the combination of the insulin-related indices on the right side of Eq (4), suggesting that $DI_{/cle}$ can be approximated as the extension of BIE to the insulin effect during glucose administration. The dimension of SIE is dL/mg, which is the same as the dimension of $1/G$. Hence, it is plausible that SIE is inversely correlated with blood glucose levels. Of note, the dimension of $k_{sec}$ in Eq (9) is 1/min, but if $k_{sec}$ is calculated based on first-phase insulin secretion, the dimension of $k_{sec}$ becomes 1. Therefore, when assessing $DI_{/cle}$ based on first-phase insulin secretion, the dimension of $k_{sen}k_{sec}/k_{cle}$ is dL/mg. This is consistent with the results shown in Appendix 1, and it is essential to consider the dimension of each parameter when determining the coefficients of $DI_{/cle}$.

##### 3. $DI_{/cle}$ and glucose effectiveness determine blood glucose levels

This appendix details how blood glucose levels are determined by $DI_{/cle}$ and glucose effectiveness. We examined a steady state during glucose administration by using a simple and stable model (7), which can be written as follows:

$$0 = -k_{glu}G - k_{sen}IG + f, \quad \text{Eq (10)}$$

$$0 = k_{sec}G - k_{cle}I, \quad \text{Eq (11)}$$

where $G$ and $I$ indicate blood glucose and insulin levels, respectively. The indices $k_{\text{glu}}$, $k_{\text{sen}}$, $k_{\text{sec}}$ and $k_{\text{cle}}$ are glucose effectiveness, insulin sensitivity, insulin secretion, and insulin clearance, respectively. $f$ is the sum of the constant baseline hepatic glucose release and the amount of infused glucose.

Eq (10) and Eq (11) can be transformed as follows:

$$f = \frac{k_{\text{sen}} k_{\text{sec}}}{k_{\text{cle}}} G^2 + k_{\text{glu}} G. \qquad \text{Eq (12)}$$

Hence, the effect of insulin on blood glucose levels may be $k_{\text{sen}} k_{\text{sec}}/k_{\text{cle}}$, taking glucose effectiveness into account. However, glucose effectiveness cannot be evaluated precisely by using this kind of simple one-compartment model (8). Further studies investigating the effect of insulin and glucose effectiveness on blood glucose levels are warranted.

One possible explanation for the difference between $\text{DI}_{\text{/cle}}$ defined in this study ($k_{\text{sen}} k_{\text{sec}}/{k_{\text{cle}}}^2$) and the analytically indicated insulin effect ($k_{\text{sen}} k_{\text{sec}}/k_{\text{cle}}$) is that the effect of insulin may depend on blood glucose levels, as shown in Appendix 2 (*i.e.*, even if insulin sensitivity, insulin secretion, and insulin clearance are the same, higher blood glucose levels can increase the insulin effect to lower blood glucose levels). Eq (12) can be transformed as follows:

$$G = \frac{f}{\frac{k_{\text{sen}} k_{\text{sec}}}{k_{\text{cle}}} G + k_{\text{glu}}}. \qquad \text{Eq (13)}$$

Blood glucose levels are affected not only by insulin sensitivity, insulin secretion, and insulin clearance, but also by glucose effectiveness; however, if blood glucose levels are approximated only by insulin clearance, Eq (8) can be experimentally indicated. Taken together, Eq (13) can be approximated and transformed as follows:

$$G \sim \frac{f}{\frac{k_{sen} k_{\text{sec}}}{{k_{\text{cle}}}^2} k + k_{\text{glu}}}, \qquad \text{Eq (14)}$$

where $k$ is the weight of DI/cle ($k_{\mathrm{sen}}k_{\mathrm{sec}}/k_{\mathrm{cle}}{}^2$) relative to glucose effectiveness ($k_{\mathrm{glu}}$) in controlling blood glucose levels. Hence, blood glucose levels during glucose administration are mainly determined by DI/cle ($k_{\mathrm{sen}}k_{\mathrm{sec}}/k_{\mathrm{cle}}{}^2$) and glucose effectiveness ($k_{\mathrm{glu}}$).

#### 4. Relation between the phenomenon of "moving up the curve" of DI/cle and a pathway of glucose homeostasis failure predicted by a previous modeling study

This appendix indicates that the phenomenon of "moving up the curve" of DI/cle for IGT (Fig. 2) represents almost the same phenomenon as predicted in the previous modeling study (9), indicating a pathway to failure of glucose homeostasis in which a change in insulin clearance alter glucose dynamics even when the normal glucose set point is maintained.

Glucose homeostasis is reported to show "dynamical compensation", a property of systems in which for every time varying input, the complete dynamics of the output are insensitive to variations in key parameters of the system (9). The glucose homeostasis circuit, which represents dynamical compensation, can be written as follows:

$$\frac{dG}{dt} = u_0 + u(t) - (k_{\mathrm{glu}} + k_{\mathrm{sen}}I)G, \qquad \text{Eq (15)}$$

$$\frac{dI}{dt} = p\beta \cdot \rho(G) - k_{\mathrm{cle}}I, \qquad \text{Eq (16)}$$

$$\frac{d\beta}{dt} = \beta \cdot h(G), \qquad \text{Eq (17)}$$

where $u_0$ and $u(t)$ are endogenous glucose production and infused glucose, respectively, $p$ and $\beta$ represent insulin secretion and beta-cell functional mass, respectively, and $\rho(G)$ is a monotonically increasing function of $G$. $h(G)$ is the beta-cell growth rate, and at the glucose set point, $G = G_0$, $h(G_0) = 0$. This system was reported to have dynamical compensation with respect to insulin sensitivity, $k_{\mathrm{sen}}$. Of note, $k_{\mathrm{cle}}$ includes both peripheral and hepatic insulin clearance, since insulin recirculates to the liver through hepatic vein and artery (10).

As previously described (9), the steady-state solution can be written as follows:

$$\frac{k_{\mathrm{sen}} p\beta \cdot \rho(G_0)}{k_{\mathrm{cle}}} = \mathrm{constant} \qquad \text{Eq (18)}$$

which can be approximated as the analytically indicated $DI_{/cle}$ is constant (Appendix 2).

The previous study (9) showed that this model does not have dynamical compensation to variation in insulin clearance, and that the change in insulin clearance alter glucose dynamics even when the normal glucose set point $G_0$ is maintained. This can be considered as a pathway for progression of glucose intolerance (9). The decrease in insulin clearance with maintained $G_0$ and with satisfying Eq (18) represent almost the same phenomenon of "moving up the hyperbolic curve" of $DI_{/cle}$. Hence, the phenomenon of "moving up the curve" of $DI_{/cle}$ at the early stage of glucose intolerance can be considered as the same phenomenon that the change in insulin clearance alter glucose dynamics even when the normal glucose set point is maintained.

There are several different methods for calculating DI, some of which are based on the product of insulin concentration and insulin sensitivity (11). However, this result indicated that an increase in insulin concentration due to increased insulin secretion and an increase in insulin concentration due to decreased insulin clearance may have different effects on glucose dynamics.

It may seem contradictory that while $DI_{/cle}$ well correlated with PG120, $DI_{/cle}$ was comparable between NGT and IGT. One possible explanation for this is that blood glucose fluctuations can be changed even with the same $DI_{/cle}$, as we described in this Appendix. Another possible explanation for this is that other kind of regulatory mechanisms such as glucose effectiveness and the effect of glucagon are related to the glucose intolerance (12,13). The other possible explanation is that there may actually be a difference in $DI_{/cle}$ between NGT and IGT because there were not enough participants to capture the statistically significant difference in $DI_{/cle}$ between NGT and IGT. Although $DI_{/cle}$ did not differ significantly between NGT and IGT, this does not mean that it was clinically equivalent in the two groups. Given our previous finding that $k_7$, a parameter for insulin clearance, was significantly reduced in individuals with IGT relative to those with NGT (4), however, the decline in $DI_{/cle}$ is not as great as that in DI in the progression from NGT to IGT. Hence, Eq (4) needs to be improved and validated in the future to reflect blood glucose fluctuations more accurately.

**5, Previously reported mathematical model cannot estimate $DI_{/cle}$ from a hyperinsulinemic-euglycemic clamp alone**

For each of the 112 subjects, we estimated the parameters of the previously reported mathematical model (4) (Fig S6A) to reproduce the time course of the hyperinsulinemic-euglycemic clamp. Some estimated parameters including $k_5$_ins and $k_7$_ins, which correspond to insulin secretion and insulin clearance, respectively, showed a tendency to diverge (*i.e.*, their value reached the predetermined upper or lower limits in some subjects) (Fig. S6B). An estimated parameter, $k_4$_ins, which corresponds to insulin sensitivity estimated from the hyperinsulinemic-euglycemic clamp alone, was statistically significantly correlated with $k_4$_full, which corresponds to insulin sensitivity estimated from both hyperglycemic and hyperinsulinemic-euglycemic clamps (Fig. S6C). However, $k_5$_ins, $k_7$_ins, and $k_5/k_7{}^2$_ins, which correspond to insulin secretion, insulin clearance, and the ratio of insulin secretion to the square of insulin clearance estimated from the hyperinsulinemic-euglycemic clamp alone, respectively, were not statistically significantly correlated with $k_5$_full, $k_7$_full, and $k_5/k_7{}^2$_full, which correspond to insulin secretion, insulin clearance, and the ratio of insulin secretion to the square of insulin clearance estimated from both the hyperglycemic clamp and the hyperinsulinemic-euglycemic clamp (Fig. S6C), respectively. Since insulin sensitivity and insulin clearance were reported to be able to be calculated from the hyperinsulinemic-euglycemic clamp (10,14), if insulin secretion or the ratio of insulin secretion to the square of insulin clearance can be estimated from the hyperinsulinemic-euglycemic clamp alone, $\mathrm{DI}_{/\mathrm{cle}}$ can be estimated from a single clamp test. Hence, we investigated whether insulin secretion or the ratio of insulin secretion to the square of insulin clearance could be estimated from the hyperinsulinemic-euglycemic clamp alone. Of note, it is neither a necessary nor a sufficient condition for estimating $\mathrm{DI}_{/\mathrm{cle}}$ that insulin secretion or the ratio of insulin secretion to the square of insulin clearance can be estimated from the hyperinsulinemic-euglycemic clamp alone; however, we examined these two to exclude the possibilities that only insulin sensitivity, which has been estimated only from the hyperinsulinemic-euglycemic clamp alone (4), can be estimated and that insulin secretion cannot be estimated at all.

Consequently, $\mathrm{DI}_{/\mathrm{cle}}$ estimated from the hyperinsulinemic-euglycemic clamp alone ($k_4 k_5/{k_7}^2$_ins) was not statistically significantly correlated with $\mathrm{DI}_{/\mathrm{cle}}$ estimated from both the hyperglycemic clamp and the hyperinsulinemic-euglycemic clamp ($k_4 k_5/{k_7}^2$_full) ($r = 0.11;\ 95\%$ CI, $-0.08$ to $0.30$) (Figs. S6C, S7A). We hypothesized that the reason for this result might be related to the model structure: in this model (Fig. S6A), glucose and insulin are injected around the pancreas ($X$ and $Y$), but they are injected in the peripheral blood during the clamp test.

#### 6, Modified mathematical model

We constructed a mathematical model of the feedback loop that links glucose and insulin (Fig. 3A) by modifying the previously reported model (4) with changing the position of influx $G$ and influx $I$, and with the addition of $k_{\mathrm{m8}}$ as follows:

$$\frac{dG}{dt} = \text{flux 1} - \text{flux 2} + \text{flux 3} - \text{flux 4} + \text{influx } G$$

$$= k_{\mathrm{m1}} Y - k_{\mathrm{m2}} G + \frac{k_{\mathrm{m3}}}{k_{\mathrm{m8}} + I} - k_{\mathrm{m4}} G I + f_1(t)$$

$$\frac{dI}{dt} = \text{flux 6} - \text{flux 7} + \text{influx } I = k_{\text{m6}}X - k_{\text{m7}}I + f_2(t)$$

$$\frac{dY}{dt} = -\text{flux 1} + \text{flux 2} = -k_{\text{m1}}Y + k_{\text{m2}}G$$

$$\frac{dX}{dt} = \text{flux 5} - \text{flux 6} = k_{\text{m5}}Y - k_{\text{m6}}X$$

where the variables $G$ and $I$ denote blood glucose and insulin concentrations, respectively. The fluxes, influx $G$ and influx $I$ denote glucose and insulin infusions, respectively.

For each subject, the parameters of the model to reproduce the time course were estimated, as described previously (4,15). In brief, each parameter of the model for plasma glucose and serum insulin concentration was estimated in the range from $10^{-4}$ to $10^{4}$. For these methods, the parameters were estimated to minimize the objective function value. The value was defined as residual sum of the square (RSS) between the measured time course during the clamp analyses and the model trajectories. RSS used in the model was given by the following equation:

$$\text{RSS} = \frac{n_I}{n_G+n_I}\sum_{i=1}^{n_G}[G(t_i) - G_{\text{sim}}(t_i)]^2 + \frac{n_G}{n_G+n_I}\sum_{i=1}^{n_I}[I(t_i) - I_{\text{sim}}(t_i)]^2,$$

where $n_G$ and $n_I$ are the total numbers of time points of plasma glucose and serum insulin, respectively, $t_i$ is the time of i-th time point. Plasma glucose and serum insulin concentrations of each subject were normalized by dividing them by the respective maximum value. $G(t)$ is the time-averaged plasma glucose level within the time range $(t - 5)$ min to $t$ min. The interval was every 1-min. $I(t)$ is the serum insulin level at $t$ min. $G_{\text{sim}}(t)$ and $I_{\text{sim}}(t)$ are simulated blood glucose and blood insulin levels, respectively. The numbers of generations and parents of the meta-evolutionary programming were 4000 and 400, respectively. Of note, identifiability of the parameters (16) of this model was not validated. Hence, even if $\text{DI}_{\text{/cle}}$ can be estimated at the population level, it may not be possible to uniquely identify all parameters in an individual.

#### 7. Prediction of DI/cle from a hyperinsulinemic-euglycemic clamp alone

This appendix details how DI/cle was estimated from a hyperinsulinemic-euglycemic clamp alone using a simple analytically computable method.

Following a formula for calculating insulin sensitivity index (ISI) (4), insulin sensitivity ($k_{\mathrm{sen}}$) can be approximated as follows:

$$0 = -k_{\mathrm{sen}} I_{\mathrm{SS}} G_{\mathrm{SS}} + f_{\mathrm{g}}, \qquad \text{Eq (19)}$$

where $I_{\mathrm{SS}}$, $G_{\mathrm{SS}}$, and $f_{\mathrm{g}}$ represent blood insulin level at the steady state during a hyperinsulinemic-euglycemic clamp, blood glucose level at the steady state during a hyperinsulinemic-euglycemic clamp, and glucose infusion rate at the steady state, respectively.

Eq (11) can be transformed as follows:

$$\frac{k_{\mathrm{sec}}}{k_{\mathrm{cle}}} = \frac{I_0}{G_0}, \qquad \text{Eq (20)}$$

where $I_0$ and $G_0$ represent blood insulin level in the fasting state and blood glucose level in the fasting state, respectively.

Following a formula for calculating metabolic clearance rate (4,5), insulin clearance ($k_{\mathrm{cle}}$) can be approximated as follows:

$$k_{\mathrm{cle}} = \frac{f_{\mathrm{i}}}{I_{\mathrm{SS}} - I_0}, \qquad \text{Eq (21)}$$

where $f_{\mathrm{i}}$ represents insulin infusion rate at the steady state. Of note, this method cannot distinguish peripheral insulin clearance from hepatic insulin clearance (10). In addition, $k_{\mathrm{cle}}$ in Eq (20) and that in Eq (21) can be different, and this method may cause some bias.

Eq (19), Eq (20) and Eq (21) can be transformed and approximated as follows:

$$\frac{k_{\mathrm{sen}} k_{\mathrm{sec}}}{{k_{\mathrm{cle}}}^2} \sim \frac{f_{\mathrm{g}} I_0 (I_{\mathrm{SS}} - I_0)}{f_{\mathrm{i}} G_{\mathrm{SS}} G_0 I_{\mathrm{SS}}} \qquad \text{Eq (22)}$$

Of note, since $k_{\mathrm{cle}}$ in Eq (20) and that in Eq (21) can be different, the values of both sides of Eq (22) are not equal.

**8. Estimation of $\mathrm{DI}_{/\mathrm{cle}}$ from an oral glucose tolerance test or a fasting blood test**

This appendix details how $\mathrm{DI}_{/\mathrm{cle}}$ was estimated from OGTT or a fasting blood test. Here, as an index of insulin secretion ($k_{\mathrm{sec}}$), we calculated the insulin secretion rate at the fasting state (3) divided by the fasting glucose level. As an index of insulin clearance ($k_{\mathrm{sen}}$), we calculated the insulin secretion rate at the fasting state (3) divided by the fasting insulin level, as previously proposed (17). As an index of insulin sensitivity during OGTT ($k_{\mathrm{sen\_OGTT}}$), we calculated oral glucose insulin sensitivity (OGIS) (18). It is worth noting that $\mathrm{DI}_{/\mathrm{cle}}$ calculated with Matsuda index (19) was relatively weakly correlated with that calculated from two clamp tests ($r = 0.28;\ 95\%\ \mathrm{CI}, 0.09$ to $0.45$). We thus used OGIS in calculating $\mathrm{DI}_{/\mathrm{cle}}$. As an index of insulin sensitivity at the fasting state ($k_{\mathrm{sen\_fast}}$), we calculated insulin sensitivity index (IS), which is the inverse of HOMA-IR (11). Of note, these insulin sensitivity-related indices may differ as to whether they are influenced by total body, peripheral, or liver insulin sensitivity, but since this has not yet been fully clarified (20), the present study examined the predictive accuracy of $\mathrm{DI}_{/\mathrm{cle}}$ without considering the type of insulin sensitivity.

$\mathrm{DI}_{/\mathrm{cle}}$ estimated from OGTT ($k_{\mathrm{sen}} k_{\mathrm{sec}} / {k_{\mathrm{cle}}}^2$_OGTT) and from a fasting blood test ($k_{\mathrm{sen}} k_{\mathrm{sec}} / {k_{\mathrm{cle}}}^2$_fast) were defined as $k_{\mathrm{sen\_OGTT}} k_{\mathrm{sec}} / {k_{\mathrm{cle}}}^2$ and $k_{\mathrm{sen\_fast}} k_{\mathrm{sec}} / {k_{\mathrm{cle}}}^2$, respectively. Because systematic errors can occur depending on which method is used to estimate $\mathrm{DI}_{/\mathrm{cle}}$, we did not apply the Bland-Altman Analysis, but only examined the correlations between each indicator.

### Supplementary Figures

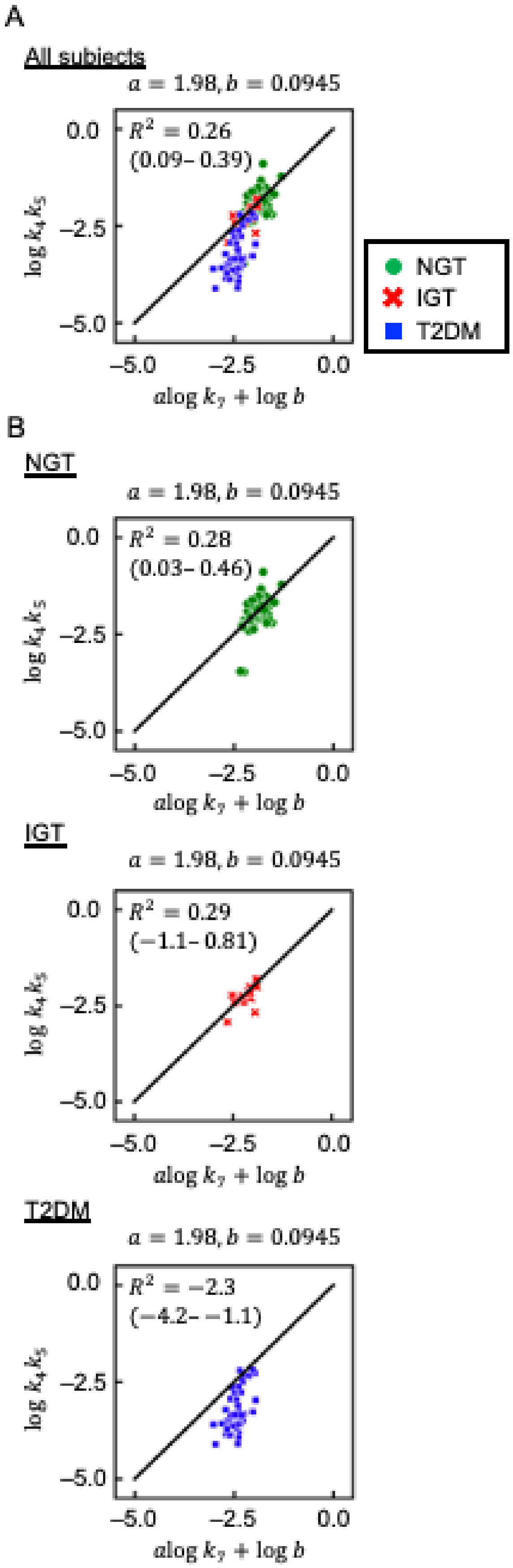

**Figure S1. Relation among insulin-related parameters.**

(A) Scatter plot for $a\log k_7 + \log b$ versus $\log k_4k_5$ (Eq (1)). Each point corresponds to the values for a single subject. Green circles, red crosses, and blue squares indicate NGT, IGT, and T2DM subjects, respectively. $R^2$ is the coefficient of determination, and the value in the parenthesis is 95% confidence interval. The parameters $a$ and $b$ were derived from the previously reported values (4), which were estimated to minimize residual sum of the square (RSS) between $bk_7{}^a$ and $k_4k_5$ in all subjects. (B) Scatter plots for $a\log k_7 + \log b$ versus $\log k_4k_5$ in NGT, IGT, and T2DM subjects (Eq (1)). The parameters $a$ and $b$ were derived from the previously reported values (4).

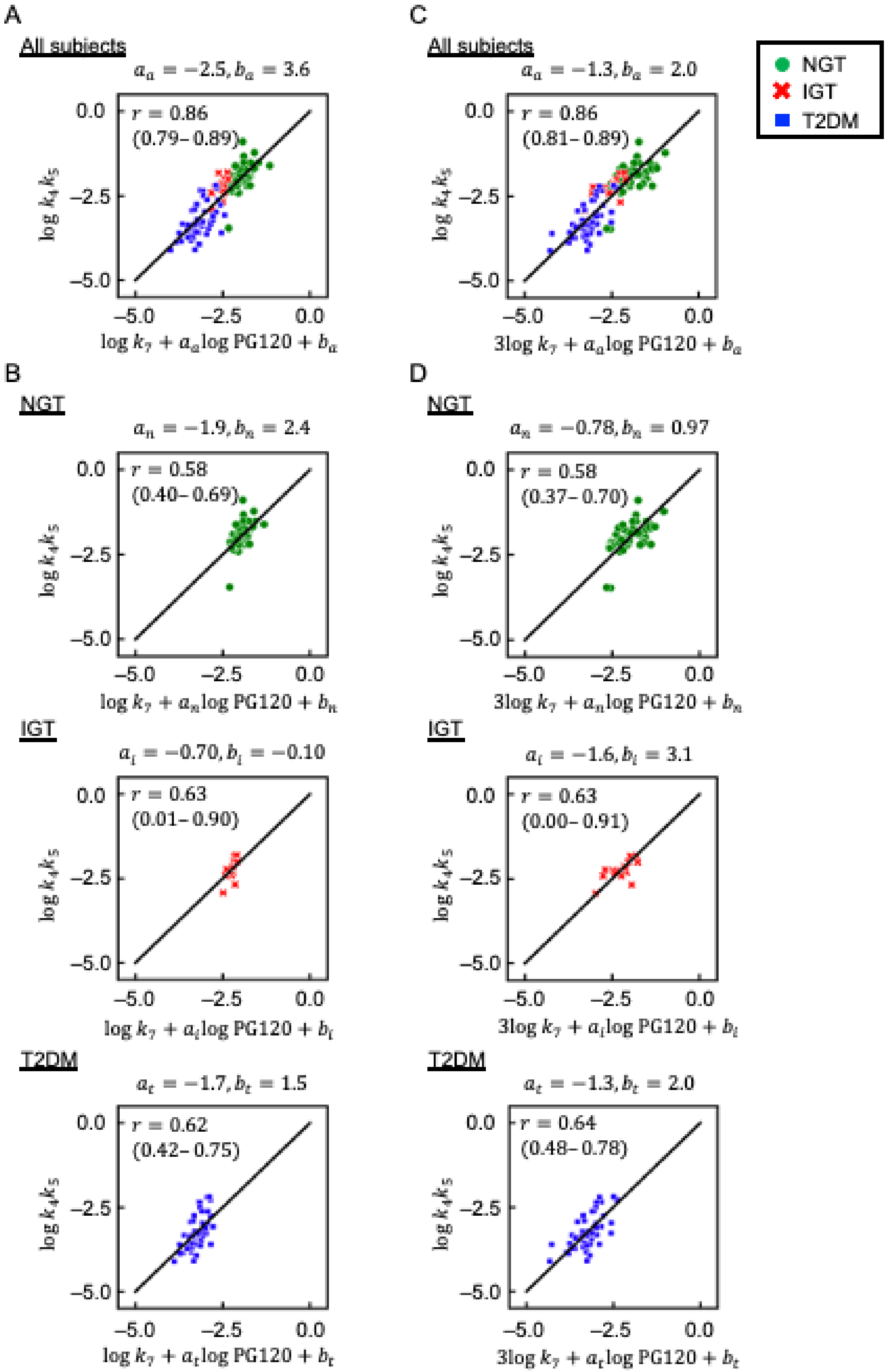

**Figure S2. Relation between insulin-related parameters and blood glucose levels.**

(A) Scatter plot for $\log k_7 + a_a \log \mathrm{PG120} + b_a$ versus $\log k_4 k_5$. Each point corresponds to the values for a single subject. Green circles, red crosses, and blue squares indicate NGT, IGT, and T2DM subjects, respectively. $r$ is Pearson's correlation coefficient, and the value in the parenthesis is 95% confidence interval. The parameters $a_a$ and $b_a$ were estimated to minimize residual sum of the square (RSS) between $\log k_7 + a_a \log \mathrm{PG120} + b_a$ and $\log k_4 k_5$ in all subjects. (B) Scatter plots for $\log k_7 + a \log \mathrm{PG120} + b$ versus $\log k_4 k_5$ in NGT, IGT, and T2DM subjects. The parameters $a_n$ and $b_n$, $a_i$ and $b_i$, and $a_t$ and $b_t$ were estimated to minimize RSS between $\log k_7 + a \log \mathrm{PG120} + b$ and $\log k_4 k_5$ in NGT, IGT, and T2DM subjects, respectively. (C) Scatter plot for $3 \log k_7 + a_a \log \mathrm{PG120} + b_a$ versus $\log k_4 k_5$. The parameters $a_a$ and $b_a$ were estimated to minimize RSS between $3 \log k_7 + a_a \log \mathrm{PG120} + b_a$ and $\log k_4 k_5$ in all subjects. (D) Scatter plots for $3 \log k_7 + a \log \mathrm{PG120} + b$ versus $\log k_4 k_5$ in NGT, IGT, and T2DM subjects. The parameters $a_n$ and $b_n$, $a_i$ and $b_i$, and $a_t$ and $b_t$ were estimated to minimize RSS between $3 \log k_7 + a \log \mathrm{PG120} + b$ and $\log k_4 k_5$ in NGT, IGT, and T2DM subjects, respectively. The indices $k_4$, $k_5$, and $k_7$, which correspond to insulin sensitivity, insulin secretion, and insulin clearance, respectively, were calculated by a previously described method (4). NGT, normal glucose tolerance; IGT, impaired glucose tolerance; T2DM, type 2 diabetes mellitus; PG120, plasma glucose concentration at 120 min during the OGTT.

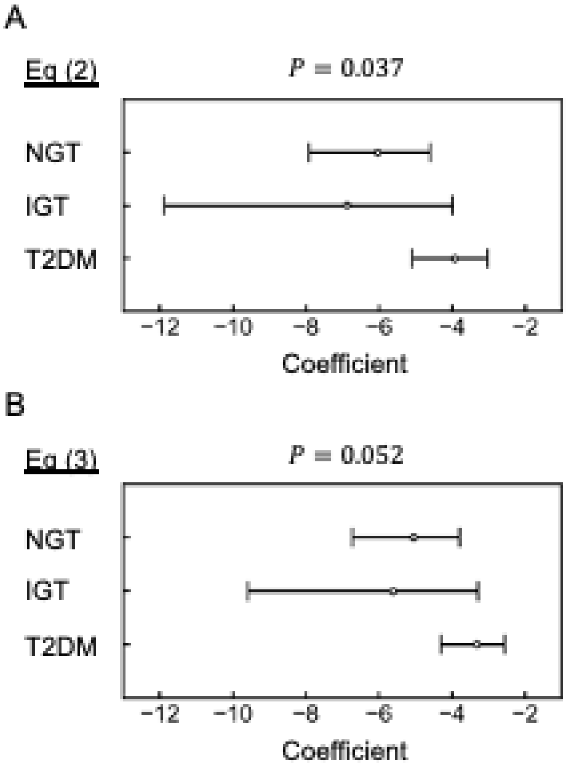

**Figure S3. Comparison of the lines estimated by Eq (2) or Eq (3) in NGT, IGT, and T2DM.**

(A) Estimated slopes of Eq (2) by the standardized major axis test. (B) Estimated slopes of Eq (3) by the standardized major axis test. Bars indicate 95% confidence intervals. The *P* value is for testing the hypothesis of no difference among groups. NGT, normal glucose tolerance; IGT, impaired glucose tolerance; T2DM, type 2 diabetes mellitus.

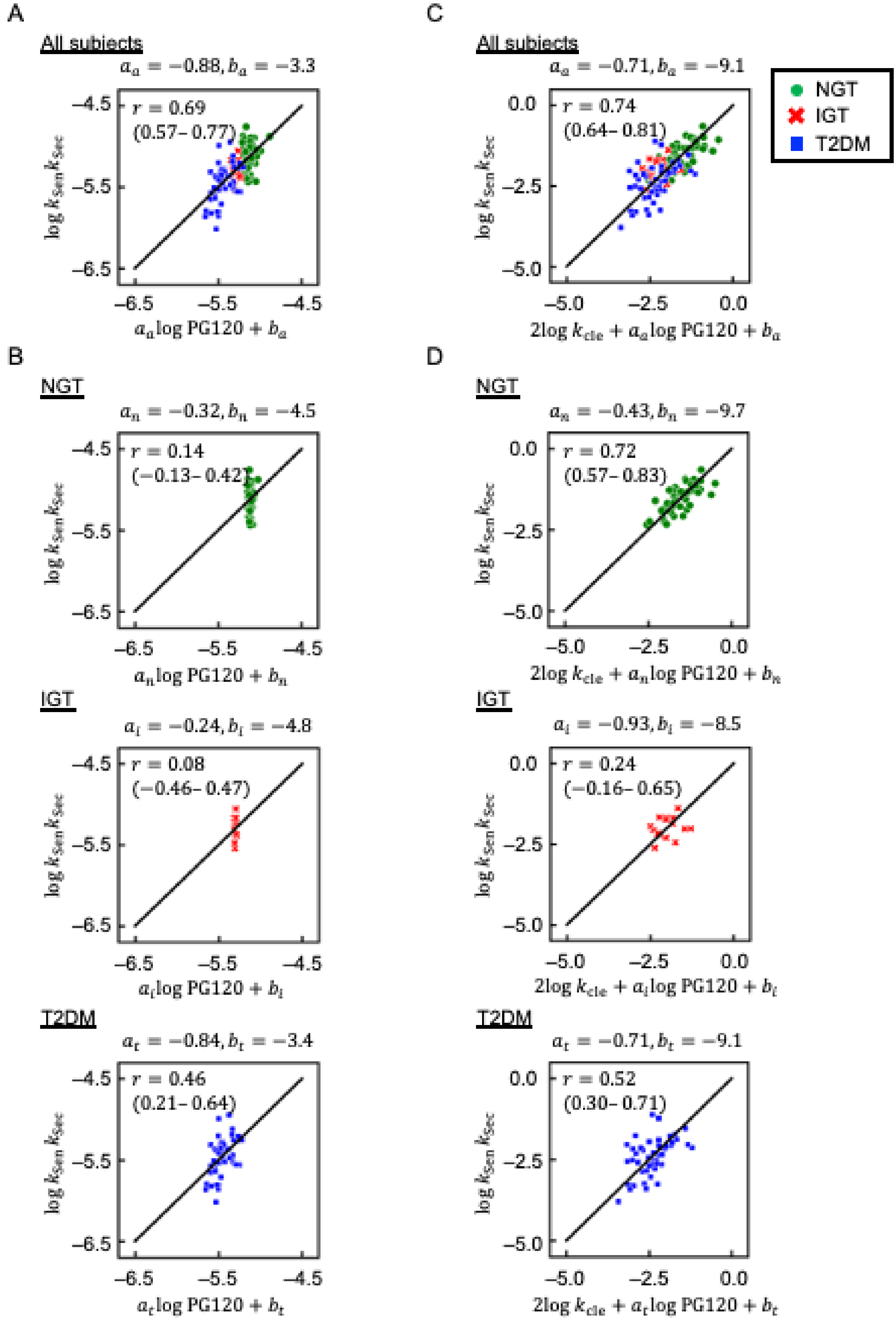

**Figure S4. Relation between insulin-related parameters and blood glucose levels.**

(A) Scatter plot for $a_a \log \mathrm{PG120} + b_a$ versus $\log k_{\mathrm{sen}} k_{\mathrm{sec}}$ (Eq (2)). Each point corresponds to the values for a single subject. Green circles, red crosses, and blue squares indicate NGT, IGT, and T2DM subjects, respectively. $r$ is Pearson's correlation coefficient, and the value in the parenthesis is 95% confidence interval. The parameters $a_a$ and $b_a$ were estimated to minimize RSS between $a_a \log \mathrm{PG120} + b_a$ and $\log k_{\mathrm{sen}} k_{\mathrm{sec}}$ in all subjects. (B) Scatter plots for $a \log \mathrm{PG120} + b$ versus $\log k_{\mathrm{sen}} k_{\mathrm{sec}}$ in NGT, IGT, and T2DM subjects (Eq (2)). The parameters $a_n$ and $b_n$, $a_i$ and $b_i$, and $a_t$ and $b_t$ were estimated to minimize RSS between $a \log \mathrm{PG120} + b$ and $\log k_{\mathrm{sen}} k_{\mathrm{sec}}$ in NGT, IGT, and T2DM subjects, respectively. (C) Scatter plot for $2 \log k_{\mathrm{cle}} + a_a \log \mathrm{PG120} + b_a$ versus $\log k_{\mathrm{sen}} k_{\mathrm{sec}}$ (Eq (3)). The parameters $a_a$ and $b_a$ were estimated to minimize RSS between $2 \log k_{\mathrm{cle}} + a_a \log \mathrm{PG120} + b_a$ and $\log k_{\mathrm{sen}} k_{\mathrm{sec}}$ in all subjects. (D) Scatter plots for $2 \log k_{\mathrm{cle}} + a \log \mathrm{PG120} + b$ versus $\log k_{\mathrm{sen}} k_{\mathrm{sec}}$ in NGT, IGT, and T2DM subjects (Eq (3)). The parameters $a_n$ and $b_n$, $a_i$ and $b_i$, and $a_t$ and $b_t$ were estimated to minimize RSS between $2 \log k_{\mathrm{cle}} + a \log \mathrm{PG120} + b$ and $\log k_{\mathrm{sen}} k_{\mathrm{sec}}$ in NGT, IGT, and T2DM subjects, respectively. The indices $k_{\mathrm{sen}}$, $k_{\mathrm{sec}}$, and $k_{\mathrm{cle}}$, correspond to insulin sensitivity, insulin secretion, and insulin clearance, respectively (Appendix 1), and PG120 is the plasma glucose concentration at 120 min during an OGTT.

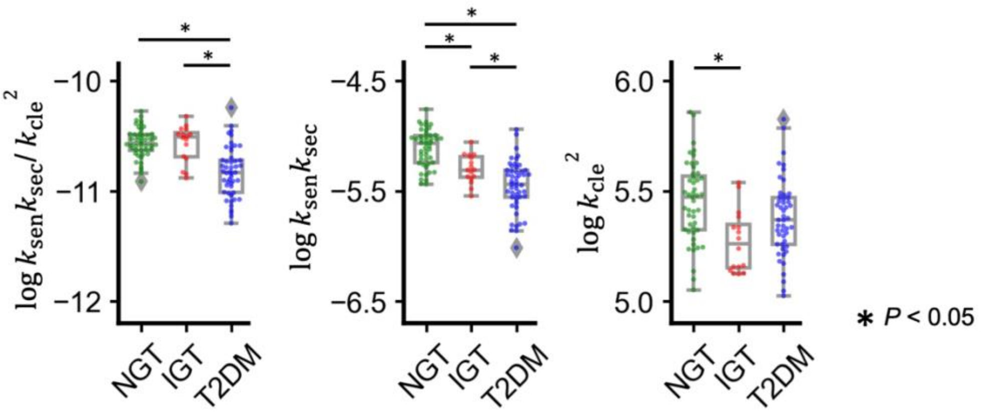

**Figure S5. DI/cle, disposition index and insulin clearance.**

Box plots of $k_{sen}k_{sec}/k_{cle}^2$, $k_{sen}k_{sec}$, and $k_{cle}^2$ in NGT, IGT, and T2DM subjects, respectively. The indices $k_{sen}$, $k_{sec}$, and $k_{cle}$ correspond to insulin sensitivity, insulin secretion, and insulin clearance (Appendix 1), respectively. The indices $k_{sen}k_{sec}/k_{cle}^2$ and $k_{sen}k_{sec}$ correspond to DI/cle and disposition index, respectively. Each point corresponds to the value for a single subject. The boxes denote the median and upper and lower quartiles. *P* values were determined by the Steel-Dwass test.

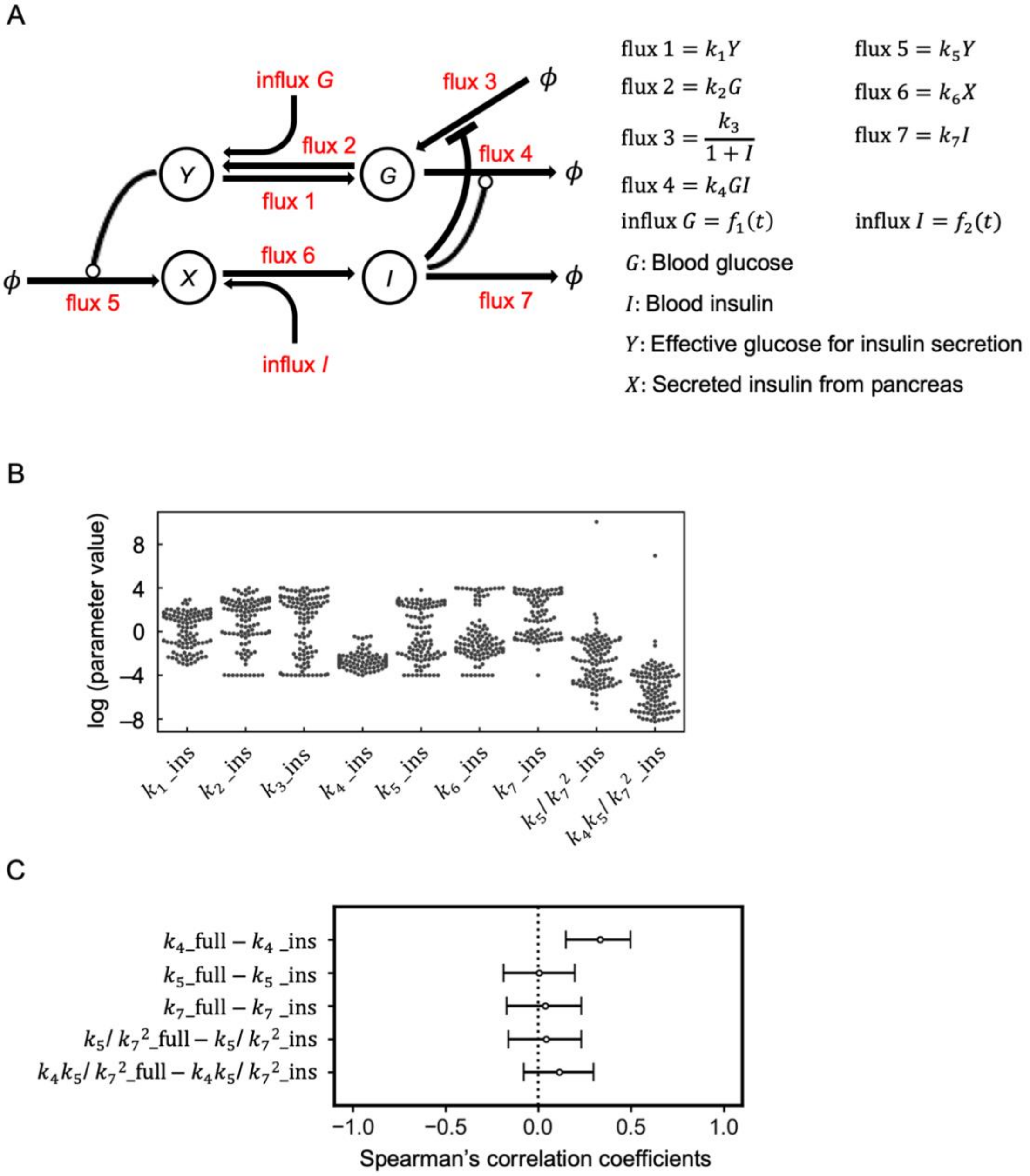

**Figure S6. Relation between parameters estimated from a hyperinsulinemic-euglycemic clamp alone and those estimated from both hyperglycemic and hyperinsulinemic-euglycemic clamps.**

(A) Model diagram of the previously described mathematical model (4) (CC-BY 4.0 license). The letters within the circle indicate the variables of the model, the arrows denote the fluxes, and the lines ending in circles or bars represent activation or inactivation, respectively. $\phi$ indicates a fixed value. The variables $G$ and $I$ represent blood glucose and insulin concentrations, respectively. The variable $Y$ corresponds to the effective glucose concentration for induction of variable $X$, which represents insulin secreted from pancreatic β-cells. (B) Estimated parameters of the previously described model (4) (Fig. S6A) based on a hyperinsulinemic-euglycemic clamp alone. The base of the logarithm is 10. (C) Spearman's correlation coefficients. Bars indicate 95% confidence intervals. The parameters $k_4$_full, $k_5$_full, $k_7$_full, $k_5/k_7{}^2$_full, and $k_4k_5/k_7{}^2$_full correspond to insulin sensitivity, insulin secretion, insulin clearance, the ratio of insulin secretion to the square of insulin clearance, and $DI_{/cle}$ estimated from both hyperglycemic and hyperinsulinemic-euglycemic clamps by the previously described mathematical model (4) (Fig. S6A). The parameters $k_4$_ins, $k_5$_ins, $k_7$_ins, $k_5/k_7{}^2$_ins, and $k_4k_5/k_7{}^2$_ins correspond to insulin sensitivity, insulin secretion, insulin clearance, the ratio of insulin secretion to the square of insulin clearance, and $DI_{/cle}$ estimated from the hyperinsulinemic-euglycemic clamp alone by the previously described mathematical model (4) (Fig. S6A).

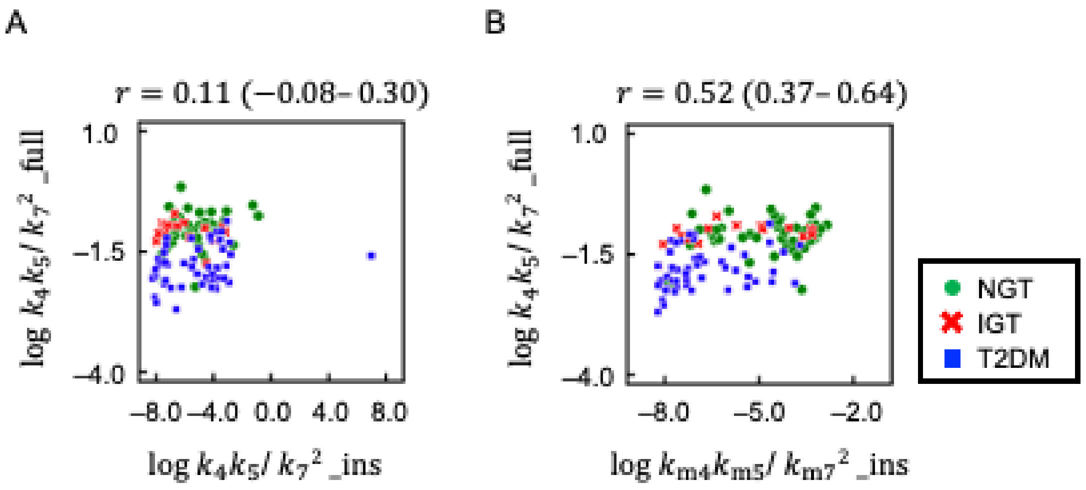

**Figure S7. Relation between DI$_{/\mathrm{cle}}$ estimated from a hyperinsulinemic-euglycemic clamp alone and that estimated from both hyperglycemic and hyperinsulinemic-euglycemic clamps.**

(A) Scatter plot for log $k_4 k_5/{k_7}^2$_ins versus log $k_4 k_5/{k_7}^2$_full. The parameters $k_4 k_5/{k_7}^2$_ins and $k_4 k_5/{k_7}^2$_full correspond to DI$_{/\mathrm{cle}}$ estimated from the hyperinsulinemic-euglycemic clamp alone and DI$_{/\mathrm{cle}}$ estimated from both hyperglycemic and hyperinsulinemic-euglycemic clamps by the previously described mathematical model (4) (Fig. S6A), respectively. Each point corresponds to the values for a single subject, with green circles, red crosses, and blue squares indicating NGT, IGT, and T2DM subjects, respectively. $r$ is Spearman's correlation coefficient, and the value in the parenthesis is 95% confidence interval. (B) Scatter plot for log $k_{\mathrm{m4}} k_{\mathrm{m5}}/{k_{\mathrm{m7}}}^2$_ins versus log $k_4 k_5/{k_7}^2$_full. The parameters $k_{\mathrm{m4}} k_{\mathrm{m5}}/{k_{\mathrm{m7}}}^2$_ins and $k_4 k_5/{k_7}^2$_full correspond to DI$_{/\mathrm{cle}}$ estimated from the hyperinsulinemic-euglycemic clamp alone by the modified model (Fig. 3A) and that estimated from the hyperglycemic and hyperinsulinemic-euglycemic clamps by the previously described mathematical model (4) (Fig. S6A), respectively. NGT, normal glucose tolerance; IGT, impaired glucose tolerance; T2DM, type 2 diabetes mellitus.

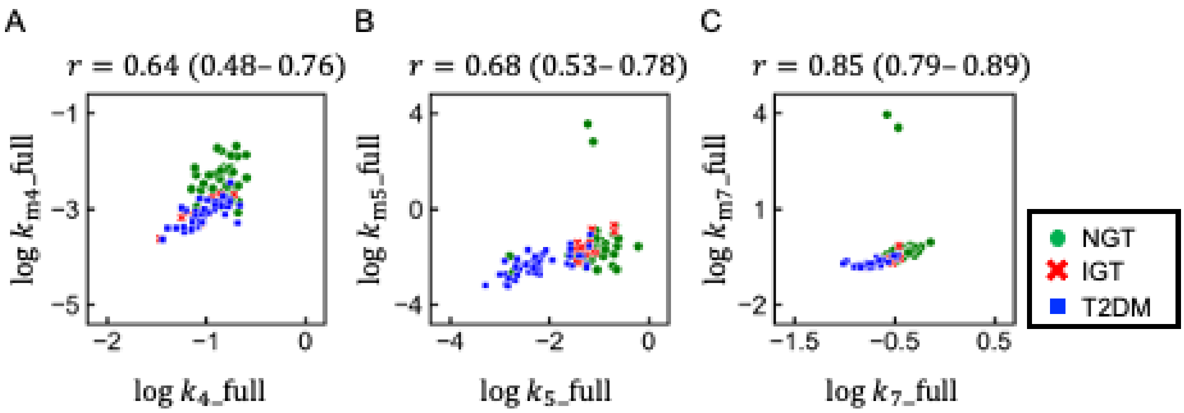

**Figure S8. Relation between parameters estimated from both hyperglycemic and hyperinsulinemic-euglycemic clamps by a previously described model and those estimated by a modified model.**

Scatter plot for log $k_4$_full versus log $k_{m4}$_full (A), log $k_5$_full versus log $k_{m5}$_full (B), and log $k_7$_full versus log $k_{m7}$_full (C). The parameters $k_4$_full, $k_5$_full and $k_7$_full correspond to insulin sensitivity, insulin secretion, and insulin clearance estimated from both hyperglycemic and hyperinsulinemic-euglycemic clamps by the previously described mathematical model (4), respectively. The parameters $k_{m4}$_full, $k_{m5}$_full and $k_{m7}$_full correspond to insulin sensitivity, insulin secretion, and insulin clearance estimated from the hyperglycemic and hyperinsulinemic-euglycemic clamps by the modified mathematical model (Fig. 3A), respectively. Each point corresponds to the values for a single subject, with green circles, red crosses, and blue squares indicating NGT, IGT, and T2DM subjects, respectively. $r$ is Spearman's correlation coefficient, and the value in the parenthesis is 95% confidence interval. NGT, normal glucose tolerance; IGT, impaired glucose tolerance; T2DM, type 2 diabetes mellitus.

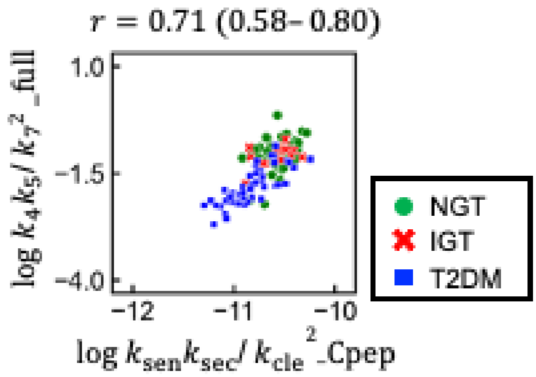

**Figure S9. Relation between DI/cle estimated from a hyperinsulinemic-euglycemic clamp alone and that estimated from both hyperglycemic and hyperinsulinemic-euglycemic clamps.**

Scatter plot for log $k_{\mathrm{sen}}k_{\mathrm{sec}}/k_{\mathrm{cle}}{}^{2}$_Cpep versus log $k_4k_5/k_7{}^{2}$_full. The parameters $k_{\mathrm{sen}}k_{\mathrm{sec}}/k_{\mathrm{cle}}{}^{2}$_Cpep and $k_4k_5/k_7{}^{2}$_full correspond to DI/cle analytically estimated from the hyperinsulinemic-euglycemic clamp alone by the serum C-peptide concentration (Appendix 1) and that estimated from both hyperglycemic and hyperinsulinemic-euglycemic clamps by the previously described mathematical model (4) (Fig. S6A), respectively. Each point corresponds to the values for a single subject, with green circles, red crosses, and blue squares indicating NGT, IGT, and T2DM subjects, respectively. $r$ is Spearman's correlation coefficient, and the value in the parenthesis is 95% confidence interval.

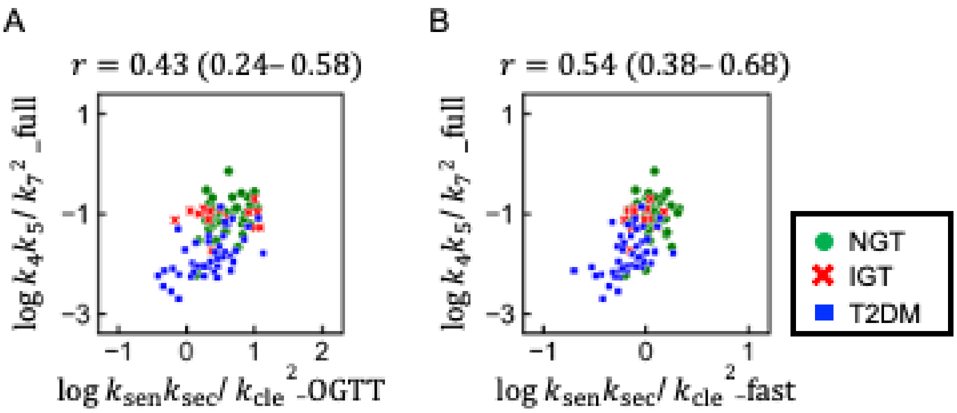

**Figure S10. Relation between $\mathrm{DI}_{/\mathrm{cle}}$ estimated from both hyperglycemic and hyperinsulinemic-euglycemic clamps and that estimated from an oral glucose tolerance test or a fasting blood test.**

Scatter plot for log $k_4k_5/k_7{}^2$_full versus log $k_{\mathrm{sen}}k_{\mathrm{sec}}/k_{\mathrm{cle}}{}^2$_OGTT (A) or log $k_{\mathrm{sen}}k_{\mathrm{sec}}/k_{\mathrm{cle}}{}^2$_fast (B). The parameters $k_4k_5/k_7{}^2$_full, $k_{\mathrm{sen}}k_{\mathrm{sec}}/k_{\mathrm{cle}}{}^2$_OGTT, and $k_{\mathrm{sen}}k_{\mathrm{sec}}/k_{\mathrm{cle}}{}^2$_fast correspond to $\mathrm{DI}_{/\mathrm{cle}}$ estimated from both hyperglycemic and hyperinsulinemic-euglycemic clamps, that estimated from OGTT, and that estimated from a fasting blood test (Appendix 8), respectively. Each point corresponds to the values for a single subject, with green circles, red crosses, and blue squares indicating NGT, IGT, and T2DM subjects, respectively. $r$ is Spearman's correlation coefficient, and the value in the parenthesis is 95% confidence interval.
